# Interparticle Crosslinked Ion-responsive Microgels for 3D and 4D (Bio)printing Applications

**DOI:** 10.1101/2025.01.28.635095

**Authors:** Vaibhav Pal, Deepak Gupta, Suihong Liu, Ilayda Namli, Syed Hasan Askari Rizvi, Yasar Ozer Yilmaz, Logan Haugh, Ethan Michael Gerhard, Ibrahim T. Ozbolat

## Abstract

Microgels offer unique advantages over bulk hydrogels due to their improved diffusion limits for oxygen and nutrients. Particularly, stimuli-responsive microgels with inherently bioactive and self-supporting properties emerge as highly promising biomaterials. This study unveils the development of interparticle-crosslinked, self-supporting, ion-responsive microgels tailored for 3D and 4D (bio)printing applications. A novel strategy was proposed to develop microgels that enabled interparticle crosslinking, eliminating the need for filler hydrogels and preserving essential microscale void spaces to support cell migration and vascularization. Additionally, these microgels possessed unique, ion-responsive shrinking behavior primarily by the Hofmeister effect, reversible upon the removal of the stimulus. Two types of microgels, spherical (µS) and random-shaped (µR), were fabricated, with µR exhibiting superior mechanical properties and higher packing density. Fabricated microgel-based constructs supported angiogenesis with tunable vessel size based on interstitial void spaces while demonstrating excellent shear-thinning and self-healing properties and high print fidelity. Various bioprinting techniques were employed and validated using these microgels, including extrusion-based, embedded, intraembedded, and aspiration-assisted bioprinting, facilitating the biofabrication of scalable constructs. Multi-material 4D printing was achieved by combining ion-responsive microgels with non-responsive microgels, enabling programmable shape transformations upon exposure to ionic solutions. Utilizing 4D printing, complex, dynamic structures were generated such as coiling filaments, grippers, and folding sheets, providing a foundation for the development of advanced tissue models and devices for regenerative medicine and soft robotics, respectively.

## 1 Introduction

Materials capable of undergoing shape transformations in response to external stimuli, such as electric field, temperature, magnetic field, light, and pH^[1]^, offer significant advantages for various applications, including biomedical implants/devices^[2–4]^, micro-mechanical and - electrical systems^[2,3]^, soft actuators^[4]^, and wearable electronics.^[5]^ Despite recent advancements, fabrication of complex shapes with programmable actuation from stimuli-responsive bioactive hydrogels remains challenging, with most examples limited to basic casting methods.^[6–8]^ Recently, 4D (bio)printing of hydrogels has emerged, integrating 3D bioprinting techniques with stimuli-responsive materials to fabricate dynamic structures capable of altering their shape or properties over time.^[6]^ Multi-material 4D (bio)printing, utilizing distinct swelling and shrinkage properties of different hydrogels, further enables anisotropic shape morphing.^[7–9]^ This process is facilitated through techniques such as extrusion-based bioprinting (EBB) with multiple (bio)inks.^[10–13]^ Among the stimuli-responsive biomaterials adapted for 4D (bio)printing, poly(N-isopropylacrylamide) (pNIPAM) and its derivates have gained widespread acceptance for the aforementioned applications.^[14–17]^ This is attributed to pNIPAM’s ability to undergo a volume phase transition when heated above its lower critical solution temperature, leading the polymer to become hydrophobic and form into a globular structure.^[18]^ Although pNIPAM exhibits adequate biocompatibility, it demonstrates poor biodegradability and lacks inherent bioactivity, requiring conjugation or functionalization with other bioactive molecules to improve its applicability in tissue engineering.^[19–21]^ Indeed, the advancement of stimuli-responsive materials that are inherently bioactive is essential to unlock tissue engineering applications.

In this direction, natural hydrogels such as gelatin, collagen, and fibrin exhibit significant potential in biomedical applications, ranging from clinical interventions to additive manufacturing.^[22]^ However, poor rheological properties of these hydrogels limit their printability, necessitating pretreatment or mixing with rheology tuning agents.^[23,24]^ Furthermore, bulk hydrogels possess nanoscale porosity^[25]^ with a low surface to volume ratio, which may not be optimal for certain processes that benefit from microscale porosity, such as cellular infiltration and neo-vessel formation during tissue regeneration and the transport of oxygen, drugs and nutrients.^[26,27]^ Thus, there is an urgent need for advanced bioinks combining printability with superior bioactive properties. In this context, microgels present a promising alternative to traditional bulk nano-porous hydrogels due to the presence of microscale void space, thereby enhancing the transport of biologics.^[28]^

Jammed microgels exhibit solid-like behavior below critical stress, known as yield stress, due to their physical interactions. Below their yield stress, densely packed microgels undergo elastic deformation.^[29]^ Above the yield stress, interparticle packing forces that resist motion are overcome, enabling the relative motion of microgels among themselves. This results in shear-thinning behavior, facilitating flow under stress and recovery after the removal of stress.^[30,31]^ This dynamic flow and recovery behavior in response to stress offers a significant advantage over conventional bulk hydrogels, aligning well with the design requirements for bioinks used in EBB.^[32]^ However, these microgels often require the incorporation of additional networks, such as a secondary hydrogel, to facilitate interparticle crosslinking, thereby binding microgels together to form stable 3D constructs suitable for handling during testing or implantation.^[12,13,33,34]^ The secondary binder hydrogel occupies interstitial spaces between microgels, consequently reducing the overall porosity of microgel-based constructs. Multiple methodologies have been documented in the literatures ^[22,28,35]^ to obviate the requirement for a binder hydrogel. For instance, dynamic covalent interparticle crosslinking was introduced, which has been demonstrated to display shear-thinning, self-healing, and shear recovery properties while maintaining cytocompatibility. Yet, these dynamic systems face challenges such as bond instability, leading to structural erosion over time.^[36]^ Furthermore, these approaches are not universally applicable to all biopolymers.^[22]^ Therefore, a versatile strategy is needed to achieve robust interparticle crosslinking for bioactive polymers while maintaining printability. Recently, 4D (bio)printing using microgel-based (bio)inks has been explored, primarily relying on pNIPAM-based polymers. These polymers undergo temperature-induced shrinking but fail to retain their shrunken state when subjected to temperature changes, highlighting a gap in the development of complex, dynamic bioprinted structures.^[13,34,37–39]^ This work addresses these challenges by advancing the design of self-supporting, bioactive microgels that eliminate the need for filler hydrogels, preserve porosity, and allow precise tuning of their stimuli-responsive behavior.

In this study, we demonstrated, for the first time, protein/carbohydrate-based self-supporting dynamic microgels that exhibit ion-responsive behavior, characterized by a reduction in volume upon exposure to ionic solutions, such as phosphate-buffered saline (PBS) due to the Hofmeister effect. These microgels were fabricated using carbohydrazide-functionalized gelatin methacrylate (FG) and oxidized alginate (OA) (**Figure 1A**). Fabrication of these microgels, which could be either spherical or randomly shaped, circumvented crosslinker-induced toxicity, as FG and OA bonded together through Schiff-base crosslinking (**Figure 1B**). Moreover, the nature of these gelatin-based microgels inherently incorporated natural cell-binding motifs, including arginyl-glycyl-aspartic acid (RGD) and matrix metalloproteases (MMPs) peptide sequences, which facilitated cell attachment and cell-mediated material resorption^[40]^, respectively, obviating the necessity for additional functionalization.^[22]^ Besides, these microgels preserved active methacrylate sites following synthesis, facilitating subsequent interparticle crosslinking via photocrosslinking (**Figure 1C**). This process obviated the necessity for infiltrating a secondary hydrogel as a binding agent, thus preserving void spaces between microgels, which was essential for cell migration and vascularization.^[41]^ Remarkably, neo-vessel networks could be developed within 7 days, demonstrated with in-ovo model, in contrast to the nearly non-recognizable capillary formation in bulk hydrogels. These mechanically stable microgels could be bioprinted to construct human-scale organs in air without any support, as demonstrated with models of human organs, such as nose and ear, and centimeter-scale perfusable devices. Furthermore, with multi-material EBB, these microgels enabled the fabrication of programmable, shape-morphing soft materials (**Figure 1D**). Specifically, the developed microgels exhibited ion-response behavior and, when co-printed with non-responsive hydrogels, formed shape-morphing constructs capable of dynamic and reversible transformations in ionic environments, governed by predefined design geometries. Likewise, this strategy, which leverages the dual crosslinking of proteins and carbohydrates to produce stimulus-responsive microgels, can be employed to other protein-based biopolymers, such as collagen and decellularized extracellular matrix (dECM), thereby broadening its applicability and advancing research in tissue engineering and regenerative medicine.

**Figure 1:**
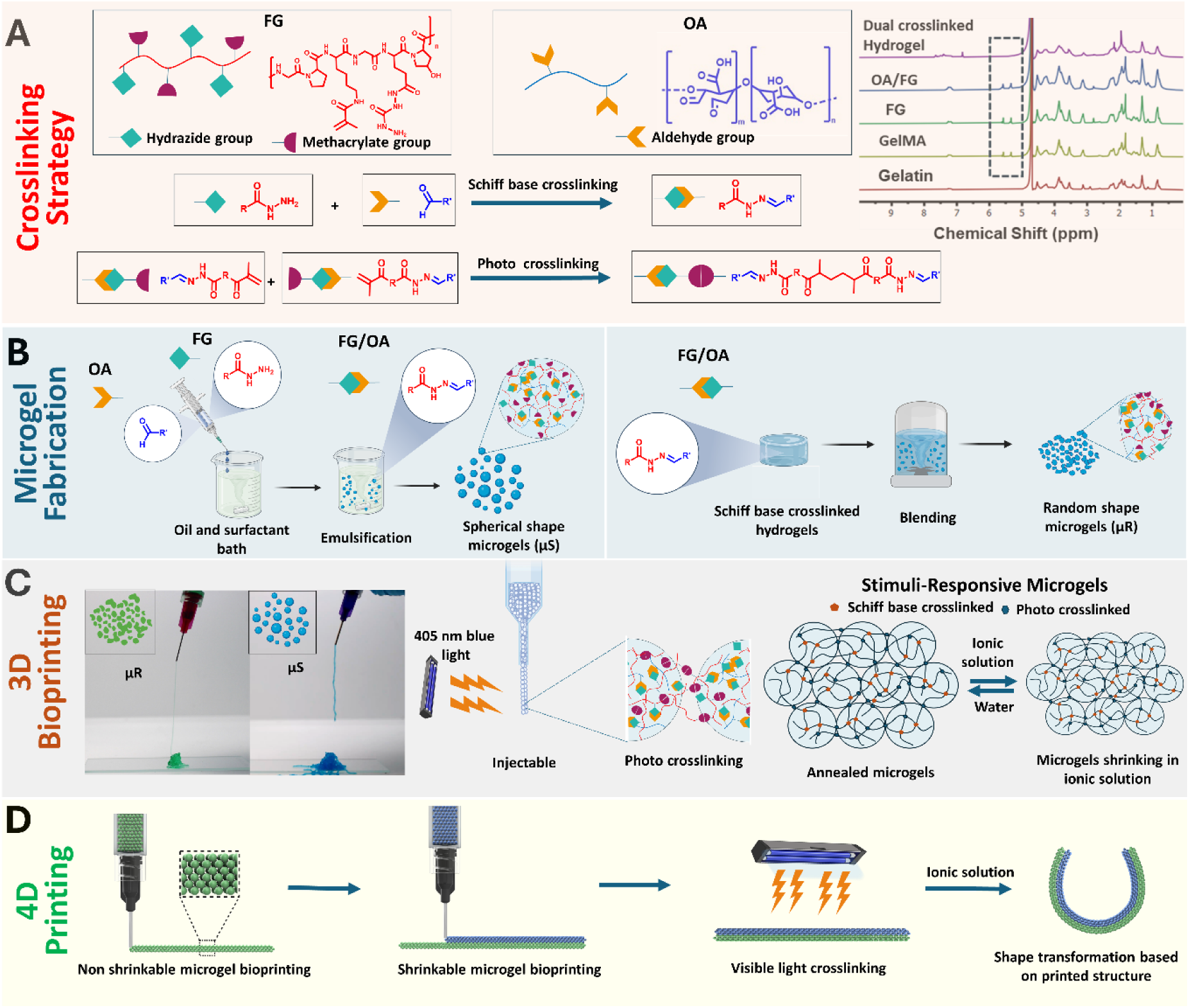
Fabrication of ion-responsive microgels. (A) Gelatin was functionalized with carbohydrazide and methacrylic anhydride to introduce hydrazide group and methacrylate groups, respectively. Alginate was oxidized to introduce aldehyde groups. NMR results confirmed the presence of peaks corresponding to gelatin functionalization, Schiff-base reaction, and photocrosslinking. (B) Multiple microgel fabrication methods, such as emulsification and blending were used to produce spherical-shaped (µS) and random-shaped microgels (µR), respectively. (C) Both types of microgels were injectable and underwent interparticle crosslinking upon exposure to 405 nm visible light. These photocrosslinked microgels were ion-responsive, showing shrinkage in ionic solution (PBS) and regaining their shape in water. (D) Multi-material 4D EBB. The first layer was printed with non-responsive gelatin methacryloyl (GelMA) microgels and hydrogel composite, followed by the printing of ion-responsive microgels. The printed construct was crosslinked with 405 nm visible light and underwent shape transformation in PBS based on the printed pattern.

## 2 Results and discussion

### 2.1 Microgel fabrication and characterization

GelMA is a widely utilized hydrogel in biomedical applications, which is synthesized from gelatin and incorporates intrinsic cellular adhesion and proliferation cues.^[42]^ It is a highly versatile material for tissue engineering, drug delivery, and 3D bioprinting due to its biocompatibility, enzymatic degradation via MMPs, cell adhesion through RGD sequences, and customizable mechanical properties.^[40,42,43]^ GelMA’s thermal stability facilitates easy tuning and functionalization, unlike other hydrogels, such as collagen and fibrinogen, which are challenging to handle and functionalize.^[44]^ Although GelMA microgels are frequently employed, bulk hydrogels are often utilized as fillers (binders) to achieve stable 3D structures.^[33,45–47]^ In certain instances, thermal and enzymatic crosslinking methods are applied to facilitate interparticle crosslinking, without the use of fillers, which may adversely affect cellular viability. To mitigate these effects, we functionalized GelMA with carbohydrazide to introduce a hydrazide group (–NH–NH_2_), a potent nucleophile that predominantly reacts with electrophilic centers, such as carbonyl groups (-C=O-) in aldehydes and ketones, and is well suited for cellular interactions owing to its biocompatibility (**Scheme S1 & S2 in Supporting Information, (SI)**).^[48]^ Coupling reactions between carbohydrazide and GelMA, resulted in FG that possessed both methacrylate and hydrazide groups (**Figure 1A**). The Schiff base reaction between the hydrazide group of FG and the aldehyde group of OA was utilized to fabricate microgels through nucleophilic reactions to form hydrazone bonds. To elucidate the kinetics of the Schiff-base crosslinking reaction, rheological analysis was conducted, measuring the storage modulus (G’) of FG/OA over time. The results indicated that G’ began to increase within 40 s, signifying the onset of the crosslinking reaction (**Figure 2A**). G’ exhibited a sharp increase for the first 5 min, followed by a gradual rise for approximately 15-20 min, after which it nearly plateaued. Compression tests were used to optimize concentrations and ratios of OA and FG, elucidating the impact of these parameters on the physical and mechanical properties of the resulting materials (**Section S1.1 & Figure S1**). Finally, two fabrication methods were used to produce µS and µR. For µS, an emulsion method was applied, wherein 5% OA and 10% FG solutions were mixed and introduced into a mineral oil and 3% Span 80 bath using a syringe (**Figure 1B**). After 30 min of crosslinking, µS were collected and washed at least three times in Milli-Q water via centrifugation. For µR, FG and OA were initially mixed to form a bulk crosslinked hydrogel, which was subsequently fragmented using a blender (**Figure 1B**). Both types of microgels retained the methacrylate group, facilitating further interparticle crosslinking, as confirmed by nuclear magnetic resonance (NMR) spectroscopy as shown in **Figure 1A**, which displayed two distinct peaks between 5-6 ppm corresponding to the methylene group in FG.^[49]^ These microgels were subsequently treated with a 5 mg/mL lithium phenyl-2,4,6-trimethylbenzoylphosphinate (LAP) solution and subjected to centrifugation. The supernatant was discarded to obtain the final microgels containing LAP. Both µR and µS demonstrated injectability (**Figure 1C**). Upon exposure to 405 nm visible light, microgels underwent interparticle photocrosslinking (**Figure 1C**). The G’ plot in **Figure 2A** showed a sharp increase upon light exposure, which validates the secondary crosslinking. The structural integrity of microgel-based constructs was validated by subjecting them to cyclic compression and relaxation tests with strains up to 20%. The results indicated that crosslinked microgels exhibited elastic deformation and complete recovery after three cycles (**Figures 2B**, **C1, C2**). In contrast, uncrosslinked microgels displayed lower stress values, attributed to the absence of interparticle crosslinking, resulting in slippage and permanent deformation leading to failure of recovery post relaxation (**Figures 2B**, **S2A-B**). Additionally, some uncrosslinked microgels adhered to the upper plate. Compression tests with up to 80 % strain or until gel failure demonstrated increasing stress with strain for crosslinked microgels (**Figures 2C3-C4**). The strain values at failure were 49 ± 1% for both crosslinked µR and µS (**Figure S2C**), while the stress at failure for µR was ∼15 kPa, significantly higher (\*\*\**p* ≤ 0.001) than that for µS (∼11 kPa) (**Figures S2D**). µR demonstrated higher stress levels compared to µS at equivalent strain levels, attributed to more efficient packing of µR (**Figures 2C1-C2**). These results validated the efficient packing and robustness of µR. Compressive moduli of crosslinked µS and µR were determined to be 62.97 ± 1.66 and 123.70 ± 4.36 Pa, respectively (**Figure 2D**). The FTIR spectra of uncrosslinked and crosslinked µR and µS are shown in **Figure S3**, highlighting reduction in the peak corresponding to double bonds at 1450 cm⁻¹, thereby confirming the occurrence of photocrosslinking.

**Figure 2:**
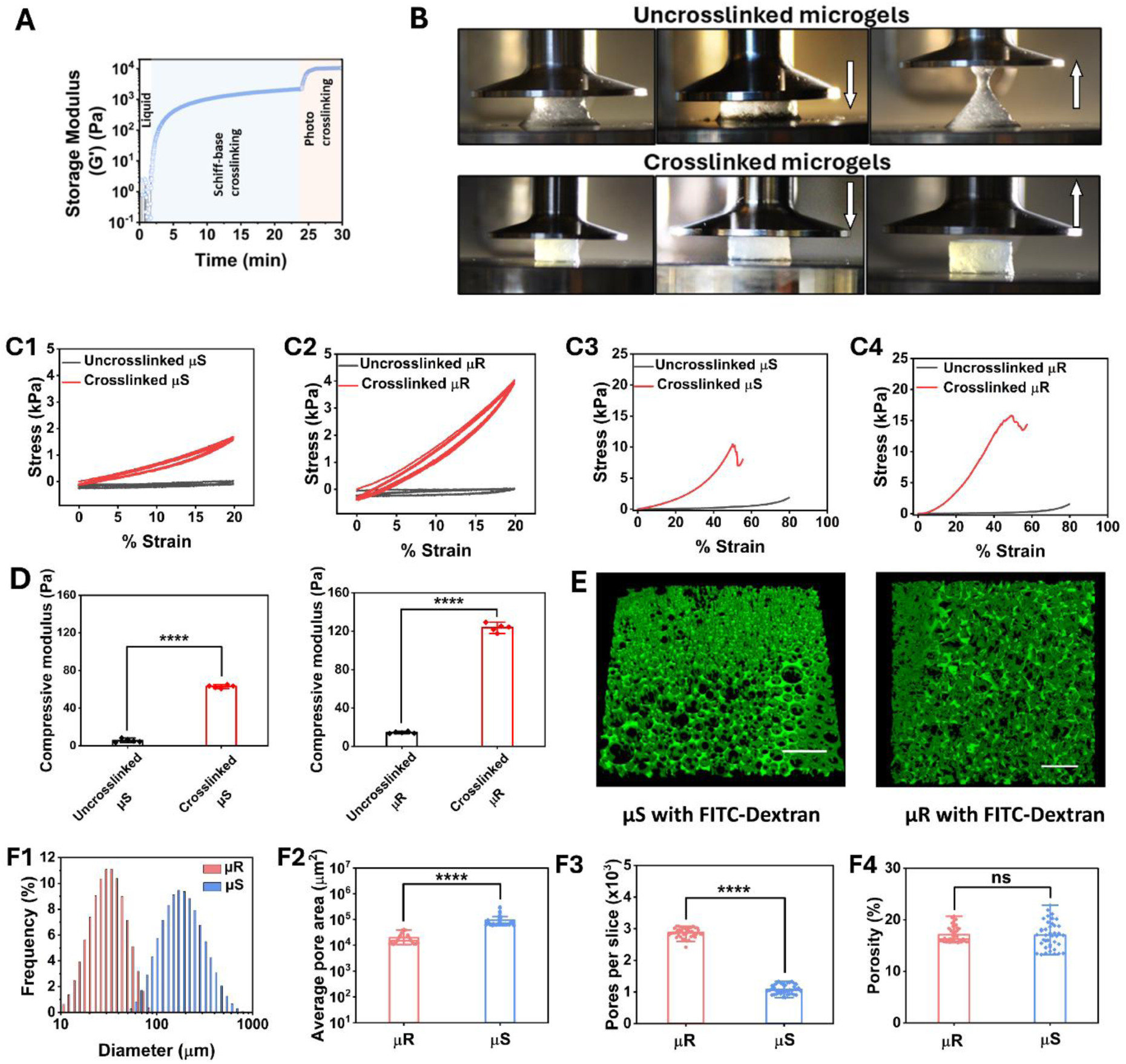
Characterization of microgels. (A) Measurement of storage modulus (G’) with time to study the kinetics of Schiff-base and photocrosslinking reactions. (B) Photos of compression testing results demonstrating slipping and breakage of uncrosslinked microgels, while validating the structural stability of interparticle-crosslinked microgels. (C1-C2) Stress-strain curves of both uncrosslinked and crosslinked µS and µR under cyclic compression and relaxation testing validating the interparticle crosslinking. (C3-C4) Stress-strain curves of both uncrosslinked and crosslinked µS and µR under compression up to 80% or failure. (D) Compressive modulus of uncrosslinked and crosslinked µS and µR showing significantly higher modulus for crosslinked microgels than uncrosslinked one (*n* = 4 independent experiments, one-way ANOVA). (E) 3D Visualization of void spaces between microgels shown with green color FITC-Dextran (scale bar: 500 µm). (F1) Microgel size distribution indicating smaller size of µR compared to µS. (F2) The average pore area of µS-based constructs was significantly higher than that for µR-based constructs. (F3) Pore count per slice showing significantly higher number of pores in µR than µS. (F4) No significant difference was found in porosity of the two different groups. Data were presented as mean ± SD where \*\*\*\**p* < 0.0001, ‘ns’ represents no significant differences.

### 2.2 Fabrication and characterization of highly porous microgel-based constructs

The packing of microgels and architecture of their pore network were visualized and analyzed. The pore network refers to the void space or pockets formed between microgels. Due to the interparticle crosslinking, these microgels did not require a secondary binder hydrogel to hold them together, which has been traditionally necessary to maintain a stable architecture. The porosity and packing density were assessed by filling the interstitial spaces with FITC-Dextran dye (2 MDa) **(Figure S4)** and analyzing z-stack images (**Figure 2E & Videos S1-S2**). Microgel size distribution analysis indicated that µR were smaller in size than µS with µS ranging from 70-750 µm and µR ranging from 10-100 µm (**Figures 2F1**). The bell curve indicates that the mean diameter of µR was ∼31 µm, whereas for µS, it was ∼163 µm. The maximum diameter observed for µR and µS as ∼100 and 750 µm, respectively. The average pore area in µR-and µS-based constructs was determined to be (0.196 ± 0.1) ×10^5^ and (0.947 ± 0.587) ×10^5^ µm^2^, respectively, with the average pore area of µS-based constructs being significantly higher (\*\*\*\**p* < 0.0001) than that of µR, owing to the large spherical shape of µS than µR (**Figure 2F2**). The pore count in µR-based constructs per plane of the z-stack image was significantly higher (\*\*\*\**p* < 0.0001) than that in µS-based constructs (**Figure 2F3**). This could be attributed to the smaller size of µR, which allowed for the incorporation of more microgels and void spaces within a given area compared to µS. Interestingly, both types of constructs did not reflect any significant difference in porosity (**Figure 2F4**). This might be attributed to µR-based constructs having a greater number of smaller pores, whereas µS-based constructs had less but larger pores. Therefore, shape and size of microgels had a direct effect on their packing, which in turn governed the number and size of pores. Further, the degradation properties of µS-and µR-based scaffolds were also studied for 28 days, and it was found both constructs were highly stable; however, µS-based scaffolds degraded more than µR-based scaffolds **(Figure S5)**.

The biocompatibility of these microgels was assessed with distinct cell types, including normal human lung fibroblasts (NHLF), human dermal fibroblasts (HDF) and GFP^+^ MDA-MB-231 cells. Bulk hydrogel (BH) was taken as control. Both µR and µS formulations demonstrated high cell viability (> 95%), indicating their excellent biocompatibility **(Figures S6, S7)**. To evaluate the impact of porosity and pore size within microgel-based constructs on neo-vessel growth, a chicken chorioallantoic membrane (CAM) assay was conducted as an in-ovo model (as illustrated in **Figure 3A**). Cylindrical constructs (with 7.57 mm in diameter, and 3.65 mm in height) made with µR, µS, and BH were implanted into eggs after 7 days of hatching, a period during which immature blood vessels form rapidly. This vascular growth continued until Day 10, reaching its terminal arrangement by Days 12-14, characterized by both sprouting and intussusceptive neo-vascular growth.^[50,51]^ During this phase, a complex vascular network, including transcapillary pillars, formed. The constructs were explanted after 14 days, where photographs revealed that microgels supported angiogenesis, with vascular networks present throughout the microgel-based constructs (**Figure 3B**). Neo-vessels successfully infiltrated void spaces between microgels and were quantified using the Angio tool as presented in **Figure S8**. The size and number of vessels were influenced by the type of microgels and their pore size. Briefly, thin vessel formation was observed in µR due to their smaller pore size, in contrast to µS, which accommodated thicker vessels. However, the control (BH) constructs were encapsulated by the embryonic tissue, with limited vascularization observed within them. This limited penetration was likely due to the hydrogel’s nanoporous nature and the absence of microporous void space. Quantification of vessel size indicated that a considerable number of vessels in µS constructs had diameters exceeding 200 µm, reaching up to 600 µm, while the vessel diameter in µR was predominantly below 150 µm (**Figure 3C**).

**Figure 3:**
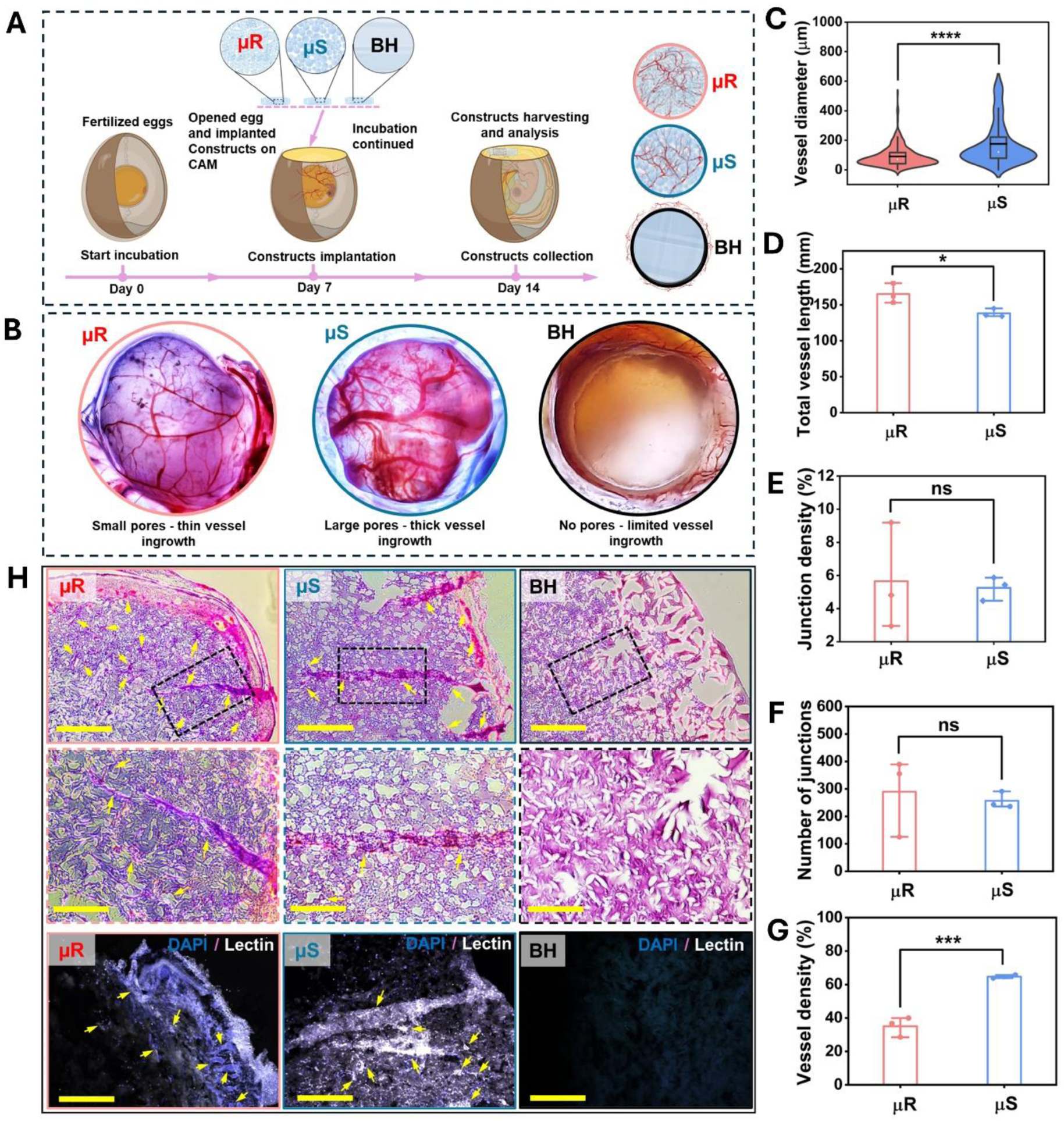
Vascularization study in-ovo. (A) Schematic of the CAM assay protocol, illustrating 7 days of egg incubation followed by construct implantation and a 7-day maturation period. (B) Photographic evidence of neo-vessel infiltration into microgel-based constructs, showing thinner vessels and capillaries in µR-based constructs, thicker vessels in µS-based constructs, and limited vessel ingrowth in BH. Quantification of vascularization showing (C) diameter of vessels, (D) combined length of all vessels infiltrating the constructs, (E) density of vessel junctions, (F) the number of vessel junctions, and (G) density of vessels. (H) H&E and DAPI/lectin-stained fluorescence images, confirming the infiltration of cells and vessels into microgel-based constructs. Data are presented as mean ± SD, *n* = 3 where *p < 0.05, ***p < 0.001 and ****p < 0.0001; ns indicates non-significant.

These results corroborated with the pore area measurements of µS and µR constructs, which suggest that vessels of up to 400 µm and 130 µm could be accommodated in µS and µR constructs, respectively (**Figure 2F2**). However, the total length of vessels was greater in µR than in µS (**Figure 3D**). Additionally, junction density and the number of junctions were slightly higher in µR compared to µS (**Figure 3E**, **F**), validating the formation of a larger number of interconnected smaller vessels in µR. The total vessel density, representing the percentage area covered by vessels, was higher in µS than in µR (**Figure 3G**) owing to the formation of thicker vessels in µS. Furthermore, histological analysis using hematoxylin and eosin (H&E) staining, as shown in **Figure 3H**, revealed successful integration of µR and µS constructs with the host tissue. Additionally, new tissue and vessels (indicated by yellow arrows) were observed in both µR and µS. In contrast, BH exhibited significantly less vessel ingrowth due to its denser microstructure lacking void space. These findings were further corroborated by fluorescent images with staining of DAPI and Lectin, which clearly indicated the presence of neo-vessels within µR and µS (yellow arrows) and the absence of vessels in BH.

### 2.3 3D (Bio)printing of microgels

3D Bioprinting with microgels offers significant potential for developing artificial tissues and organs. The inherent microporosity of microgels facilitates efficient cell loading and promotes cell migration, vascularization, and nutrient transport.^[52,53]^ To investigate the printability of these microgel-based bioinks, their rheological properties were evaluated as presented in **Figures 4A1-A4**. Shear ramp testing demonstrated that all microgels exhibited shear-thinning behavior, as evidenced by a decrease in viscosity with increasing shear rates (**Figure 4A1**), confirming their extrudability and printability for EBB. Shear recovery behavior exhibited remarkable self-healing properties (**Figure 4A2**), achieving full recovery upon the removal of stress. Such characteristics are crucial for EBB, combining extrudability upon application of the shear force followed by rapid solidification upon stress removal. Additionally, microgels fluidize under stress when a needle passes through them, functioning effectively as a support bath (**Figure 4A3**). Amplitude sweep testing results (**Figure 4A3**) indicated that G’ exceeded G’’, confirming that microgels existed in an elastic solid state at low stress levels. Upon reaching a specific stress threshold, microgels transitioned to a yielding state, indicated by a decrease in G’ and an increase in G”. At the crossover point, where G’ is equal to G’’, materials transitioned from solid to liquid phase. Beyond this crossover point, the materials exhibited liquid-like behavior, validating their yield stress properties and facilitating smooth extrusion through the nozzle as well as unobstructive movement of the nozzle through the gel without creating any crevices. The stress and strain at the crossover point were considered as the yield stress (σ) and yield strain (γ), which were found to be 64.28 ± 6.69 kPa and 13.86 ± 1.40%, respectively, for µR. These values were notably greater (\*\*\**p* < 0.001) than those for µS, with σ of 15.88 ± 0.38 kPa and γ of 3.95 ± 0.36% (**Figure 4A4**).

**Figure 4:**
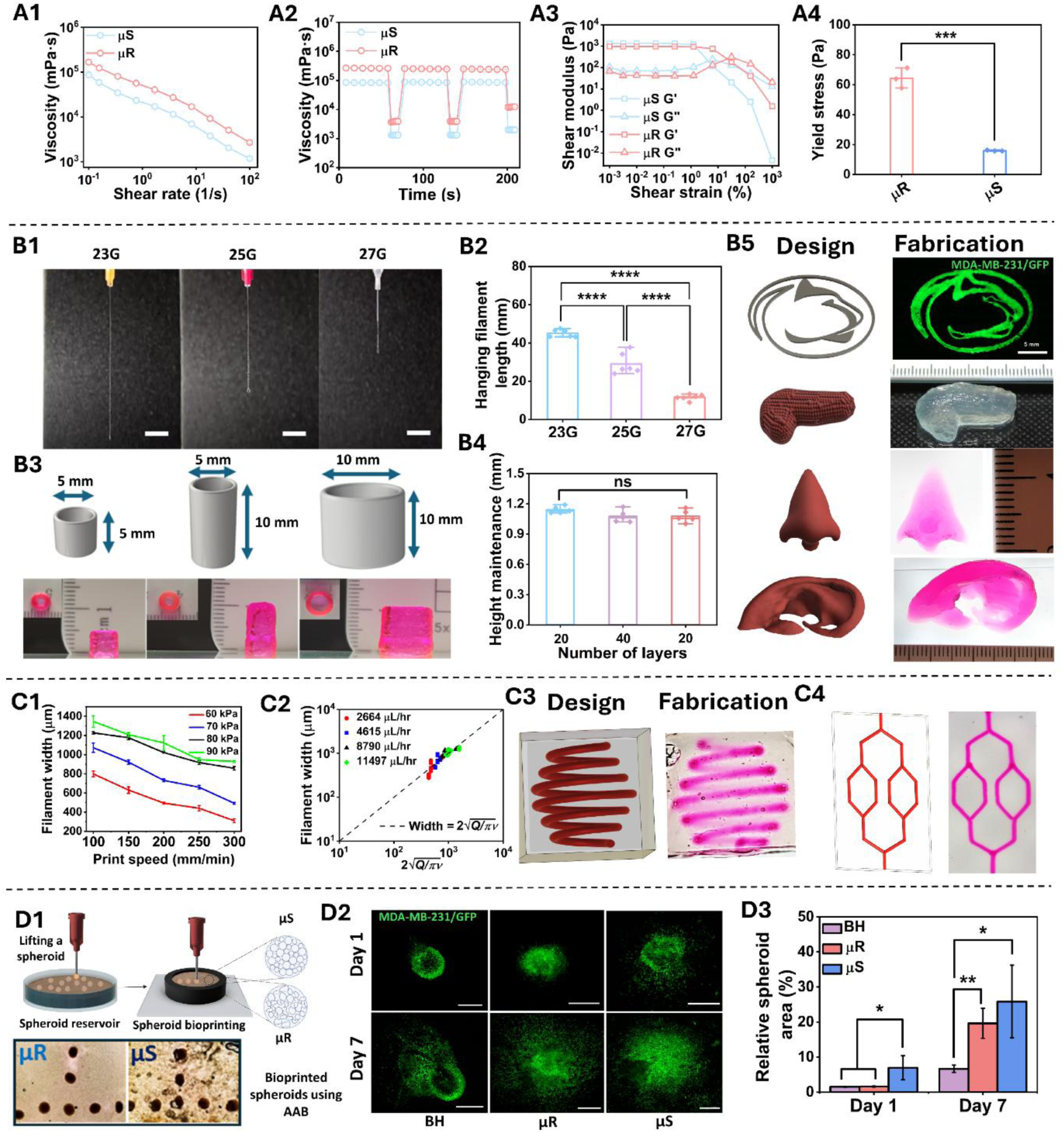
3D (Bio)printing with microgels. Rheological characterization demonstrating: (A1) shear-thinning property, (A2) self-healing capability, (A3) amplitude sweep curves for yield stress property, and (A4) yield stress values for µS and µR (*n* = 3 independent experiments, one-way ANOVA). EBB: (B1) filament hanging test demonstrating the formation of uniform µR filaments using various nozzles (23G, 25G, and 27G), (B2) quantitative analysis of hanging filament tests for 23G, 25G, and 27G nozzles, respectively (*n* = 6 independent experiments, one-way ANOVA), (B3) fabrication of scalable 3D hollow cylindrical constructs of varying dimensions using µR, (B4) validation of height maintenance in cylindrical constructs (*n* = 4 independent experiments, one-way ANOVA), (B5) bioprinting of the Penn State Nittany Lion logo with µR loaded with GFP^+^MDA-MB-231 cells. Demonstration of high-fidelity printing of human-scale organ models, such as the pancreas, nose, and ear, validating the scalability and complexity of constructs. Embedded bioprinting: (C1) variation of sacrificial filament (composed of 1% XG) width with respect to print speed and applied pneumatic pressure in the µR support bath, (C2) comparison of experimental and theoretical values of filament width, (C3) 3D spiral and (C4) a branched vasculature achieved via embedded printing of the sacrificial ink in the µR support bath, followed by perfusion with Rhodamine B dye. Aspiration-assisted bioprinting: (D1) bioprinting of HDF/GFP^+^ MDA-MB-231spheroids in µR and µS support baths, (D2) confocal images showing spheroid spreading after Day 1 and 7 (scale bar – 500 µm), (D3) quantitative assessment of spheroid spreading, demonstrating significantly higher spreading in microgels compared to the bulk hydrogel. Data were presented as mean ± SD, where ‘ns’ indicates non-significant, *p < 0.05, **p < 0.01, ***p < 0.001 and ****p < 0.0001.

Rheological assessment revealed that µS exhibited low yield stress, depicting possible non-uniform material extrusion under constant pressure. Preliminary extrusion observation indicated unstable filament formation during extrusion through a nozzle (>20G), and frequent occurrence of nozzle clogging. In contrast, high yield stress of µR facilitated its successful application in bioprinting scalable constructs. Consequently, µR was deemed more suitable for EBB applications and was utilized in EBB experiments for the rest of the study.

To evaluate the suitability of µR as a bioink, a hanging filament test was conducted, demonstrating that µR could form straight filaments with reproducible lengths of 45.10 ± 1.95 mm, 29.07 ± 5.53 mm, and 11.66 ± 1.52 mm from 23G, 25G, and 27G nozzles, respectively. The filament uniformity test indicated promising results, as continuous extrusion of µR was achieved with 23G, 25G, and 27G nozzles without any observed discontinuity or breakage (**Figures 4B1-B2**). Printability and filament fusion tests, conducted by printing grid structures with increasing grid size from 1 mm × 1 mm to 5 mm × 5 mm, demonstrated high print fidelity of µR. Edges of the smallest grid (1 mm × 1 mm) merged, resulting in the absence of pore formation and achieving 100% diffusion. The diffusion percentage decreased with an increase in grid size, reducing to 21% for a 5 mm × 5 mm grid (**Figure S9A**). The quantitative analysis of printing fidelity suggested that µR exhibited appealing printability with *Pr* ∼1 (**Figure S9B**). A filament collapse test was conducted to evaluate the stability of a horizontally suspended filament between two points due to gravity (**Figure S10**). The results demonstrated that µR filaments remained suspended without collapsing with an increase in the collapse area factor corresponding to an increase in the pillar distance (**Figure S9C**). These characterizations validated the significant potential of µR as a bioink for EBB applications. The capability of µR as a bioink for fabrication of scalable constructs was validated via printing hollow cylindrical towers with dimensions of 5 mm × 5 mm (diameter × height) (20 layers), 5 mm × 10 mm (40 layers), and 10 mm × 10 mm (40 layers). These large constructs were successfully printed with consistent height maintenance, as depicted in **Figures 4B3-B4**. Next, a 2.5D model of Penn State Nittany Lion logo and a grid was successfully bioprinted with µR loaded with GFP^+^ MDA-MB-231 cells (**Figure 4B5, S9D1-D4**), exhibiting homogeneous distribution of cells within the construct. Furthermore, the capability of microgel bioink (µR) for the construction of anatomically relative structures was validated through the fabrication of human nose and ear (**Figure 4 B5 & Video S3**). This demonstrates the potential of µR for fabricating volumetric complex structures with high shape fidelity.

The versatility of µR enables scalable bioprinting while offering the potential for integration of embedded bioprinting to create perfusable, vascularized constructs.^[54,55]^ To evaluate the suitability of µR as a support bath for embedded bioprinting, filaments composed of 1% xanthan gum (XG) were printed at varying speed and flow rates using a 25G nozzle within an µR-based support bath. The filament width decreased with increasing printing speed or decreasing flow rates (**Figures 4C1, S11**). The filament width was further compared with values predicted by the fluid continuity equation, revealing that experimental results closely matched with theoretical predictions (**Figure 4C2**). Additionally, a 3D spiral was generated with XG within µR-based support bath, followed by its removal, resulting in a perfusable 3D spiral channel within µR, which was perfused with Rhodamine B dye (**Figure 4C3)**. Similarly, microfluidic channels were printed using 1% XG within µR-based support bath loaded with GFP^+^ MDA-MB-231 cells, which was subsequently perfused with Rhodamine B dye (**Figure 4C4**). The dye began diffusing within a few minutes, as shown in **Figure S12**. To illustrate an anatomically-relevant organ model, a 2.5 cm long pancreas, along with associated vasculature and bile ducts, was printed using the intra-embedded bioprinting technique (**Figures S13**) reported recently^[56]^. To achieve this, simultaneous extrusion and support bath functionalities of µR were utilized. First, embedded printing of µR was performed within a sacrificial support bath composed of 1% XG to develop a 3D pancreatic model. This was followed by intra-embedded printing of the vasculature using XG within the printed pancreatic model, thereby creating a complex 3D structure that mimics the human pancreas. This demonstrates the versatility of µR, as it can be extruded in a free-form manner within a sacrificial bath to achieve complex 3D shapes as well as utilized as a functional support bath for 3D printing of perfusable pre-vascularized tissue models and devices.

To further demonstrate the utility of microgels, spheroid bioprinting was conducted within microgel bath utilizing aspiration-assisted bioprinting (AAB) ^[57–59]^. Human dermal fibroblasts (HDF)/GFP^+^MDA-BM-231 spheroids were prepared and bioprinted in both µR and µS support bath as shown in **Figure 4D1,** as we proposed previously.^[57–59]^ The accuracy of bioprinting was assessed, revealing that bioprinting accuracy improved with decreasing microgel size, as indicated by a significantly higher accuracy for µR compared to µS (**Figures S14**). Additionally, cell migration and spreading of spheroids within the microgel bath (µR or µS) were investigated after incubation for Days 1 and 7 (**Figure 4D2)**. The spreading area of cells from spheroids was obviously higher in microgel-based scaffolds compared to the BH control group due to highly porous structures of microgel-based scaffolds. The quantitative analysis of the coverage area of cell migration and spreading at Days 1 and 7 (**Figure 4D3**) further validated our findings. Migration area of cells within microgel baths was significantly higher than the BH group and further increased over time. Overall, fabricated bioactive microgels exhibited exceptional versatility and reliability as a support bath, enabling the fabrication of complex, anatomically accurate, and vascularized tissue constructs. These properties position the bioactive microgels as a promising material for advancing bioprinting applications.

### 2.4 4D Printing of pre-fabricated microgels

Based on the promising and intriguing findings of µR and µS in 3D (bio)printing, we further explored their potential for 4D printing. Firstly, we found an intriguing phenomenon that the bulk hydrogel and microgels synthesized in deionized (DI) water exhibited shrinkage when exposed to PBS and subsequently regained their original shapes upon re-immersion in DI water (**Figure 5A & Video S4-S5**). The schematic and microscopic images depicting the shrinking of individual µS in the presence of PBS were presented in **Figure 5A**. Then, the absolute diameter change of individual µS were measured and their shrinkage was plotted in **Figure 5B1**. The final shrinkage (%) of microgel diameter, area, and volume in PBS were 71.49 ± 6.09%, 51.47 ± 8.79% and 37.32 ± 9.58%, respectively (**Figure 5B2**). Furthermore, the shrinkage of microgel-based constructs was evaluated, revealing a 15% reduction in the size of the fabricated constructs prepared by casting (**Figures 5C1-C2**). These data were utilized to calibrate a simulation model (**Figure S15A**). A comparative analysis of the shrinking kinetics of µR-, µS-based constructs and BH over 275 s indicated that the shrinkage rate of microgel-based constructs was faster than that of BH, due to the availability of void spaces in microgel-based constructs, facilitating more rapid ion diffusion compared to the solid BH disk (**Figure 5C3 & Video S5**). The decrease and subsequent increase in the constructs area upon immersion in PBS and water, respectively, were illustrated in **Figure 5C4**. µS-based constructs shrank and regained their shape within 30 min, while µR-based constructs and BH required 40 and 53 min, respectively.

**Figure 5:**
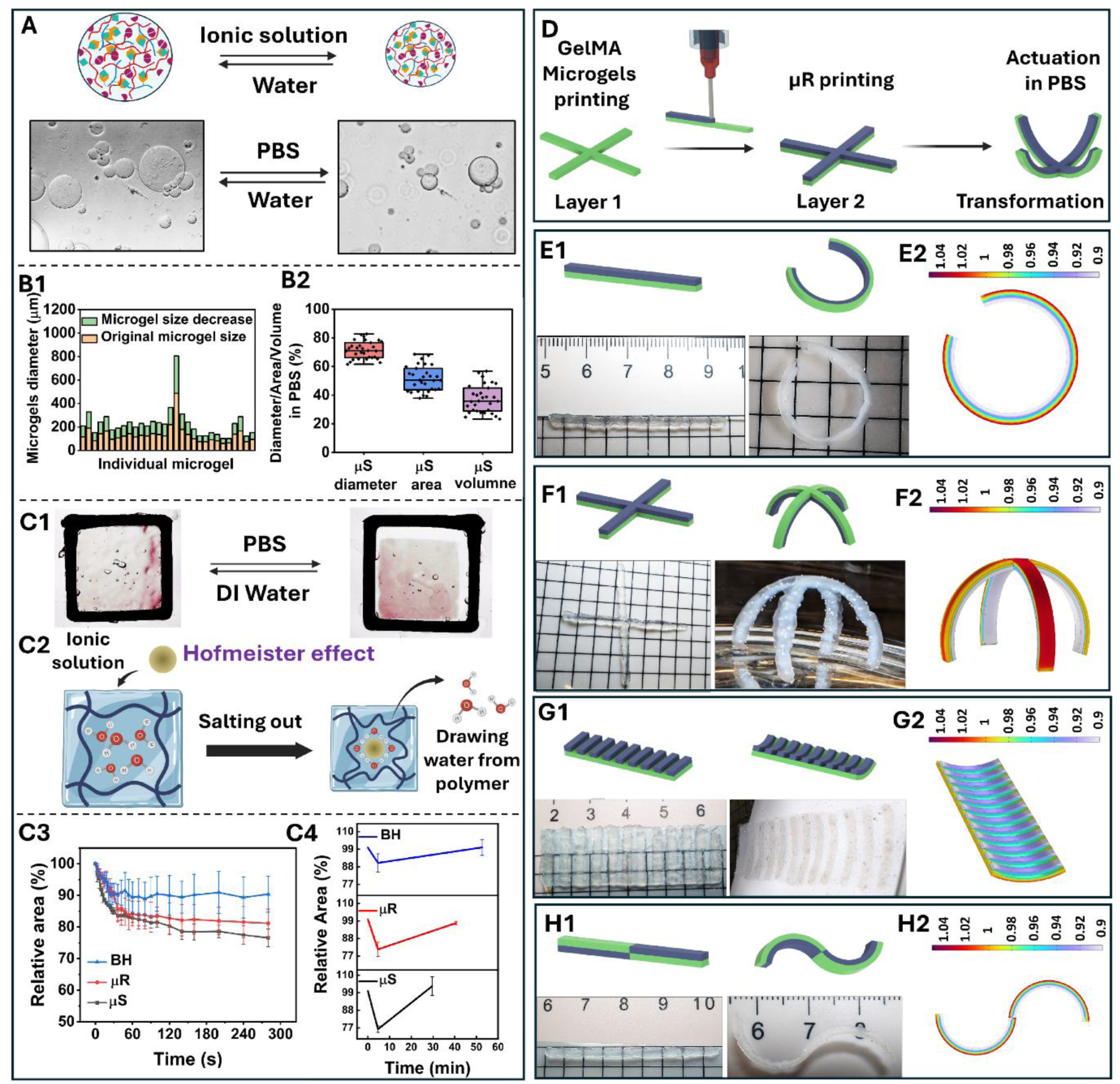
4D Printing with ion-responsive microgels. (A) Schematic and micrographs demonstrating the reversible shrinkage of individual µS. (B1) Decrease in individual microgel diameter in PBS, (B2) relative diameter, area and volume after immersion in PBS. Shrinking of microgel-based constructs: (C1) reversible shrinkage of µR-based constructs, (C2) mechanism of shrinkage based on the Hofmeister effect, (C3) shrinkage kinetics of µR-and µS-based constructs, and BH in PBS studied for 275 s (*n* = 3 independent experiments), (C4) shrinkage in PBS and subsequent shape recovery in DI water for microgels and BH. (D) Multi-material layer-wise 4D printing for shape transformation upon stimulation. Pattern-based shape transformation: (E1) a bi-layered straight line segment coiling into a circular shape and (E2) corroboration with simulation results where stretch ratio represented as heat map, (F1) a cross-shaped bi-layered structure transforming into a gripper and (F2) corroboration with simulation results, (G1) a flat structure having horizontal lines exhibiting sheet folding and (G2) corroboration with simulation results, (H1, H2) transformation of a straight filament into an S-shape upon stimulation, as predicted by simulation. Data were presented as mean ± SD.

Additionally, microgel-based constructs were also treated with different ionic (LiCl, NaCl, KCl, CaCl_2_ and MgCl_2_) and non-ionic solutions (DI water, sucrose and dimethyl sulfoxide) to understand and validate the mechanism of shrinking in the presence of ions **(Video S6-S7).** The results showed that the order of shrinkage was Ca²⁺ > Mg²⁺ > K⁺ > Na⁺ > Li⁺. The observed shrinkage behavior could be attributed to the Hofmeister ion effect, which describes the differential ability of ions to influence the arrangement as well as structure of water molecules within the polymer network **(Figure 5C2)**.^[60–62]^ Among the monovalent ions, the order of shrinkage (K⁺ > Na⁺ > Li⁺) suggested unique ion-specific effects related to their hydration properties and interactions with the polymer. K^+^, Na⁺ and Li⁺ are more kosmotropic ions and tend to get strongly hydrated. Therefore, they drew water molecules away from the polymer and induced stronger salting-out effect by breaking the hydration shell around polymeric chains,^[63]^ which resulted in polymer precipitation or shrinking in the hydrogel system. In our experiments, Ca²⁺ and Mg²⁺ induced the most significant shrinkage, likely due to their high charge density. This high charge density not only affected the structuring of water molecules but also enhanced ionic interactions with OA present in microgels promoting ionic crosslinking. Consequently, this led to the highest contraction of polymer chains by facilitating water expulsion. These findings suggested that shrinkage in this system was governed not only by the Hofmeister effect but also by specific ion-polymer interactions and the structural characteristics of the polymer network. In contrast, GelMA microgels, as a control group, did not exhibit any shrinkage in PBS (**Video S8**). To elucidate the effects of temperature, concentration and pH on the shrinkage process, experiments were conducted using 1X PBS at pH 1 and 14 and temperatures of 4 and 37 °C. Intriguingly, these experiments revealed no noticeable impact of pH or temperature on the shrinkage behavior **(Videos S9)**. However, higher concentration, where experiment was conducted using 10X PBS, exhibited faster shrinkage kinetics (**Video S10**). Together these results suggest that shrinkage was primarily driven by ion diffusion, which could be enhanced by increasing construct porosity or ion concentration.

Based on these results above, multi-material constructs incorporating ion-responsive and non-responsive microgels were co-printed to achieve predicted configurations based on their designs. As shown in **Figure 5D**, the initial layer was printed with non-responsive GelMA microgels loaded with GelMA hydrogel, then the second layer was printed with µR to form a dual-layer cross-shaped structure. Besides, various patterns were designed and printed to demonstrate programmable changes in the structure, showcasing the method’s 4D capabilities. Exemplarily, a straight filament of GelMA microgels, 8.5 cm in length, was printed, followed by a second layer of µR. The layers were crosslinked with 405 nm visible light post printing. Upon immersion in 1X PBS, the straight filament coiled into a circular shape due to the shrinkage of the upper layer, while the GelMA microgel layer remained unchanged (**Figure 5E1**). The shape-changing process was validated through physics-based modeling, showing complete agreement with experimental results (**Figure 5E2**). The simulation results indicated that the GelMA microgel layer experienced stretching, while the µR layer underwent shrinkage. The stress profile within the two materials due to shrinkage was depicted in **Figures S15B-E**, with the interface experiencing a maximum stress of 4kPa, decreasing towards the boundary. Moreover, more complex designs were also printed to validate the scalability of this approach. The cross-shaped structure with 3.8 cm long bi-layered filaments transformed into a gripper upon immersion in PBS (**Figure 5F1**); and a rectangular structure (length × width, 4.8 cm × 1.5 cm) with 12 lines of µR printed at equal distance (0.25 cm) exhibited sheet folding behavior upon stimulation (**Figure 5G1**). The simulation results corroborated with experimental observations (**Figures 5F2 and G2**), validating control over the process and confirming the accuracy of the predicted structures. To further explore its flexibility, an S-shaped central symmetrical structure (**Figure 5H1**) was designed and modeled to predict its transformation. The printed design, in which half of the printed part was composed of ion-responsive material above and below the non-responsive material, transformed into an S-shape as the ion-responsive material coiled upwards and downwards (**Figures 5H1-H2**).

In tissue engineering, there is growing interest in developing tubular and blood vessel-like structures, such as vascular networks and kidney proximal tubules.^[64–67]^ Numerous strategies have been explored to fabricate and modify printed channels, with the goal of achieving smaller dimensions to mimic smaller vessels.^[68–70]^ In this regard, we further explored our 4D printing approach to show that highly porous microgels can dynamically reduce the size of printed channels. The results showed ∼35% decrease in diameter of channels created in µR-based scaffolds with 10X PBS at room temperature **(Figure S16 & Video S11).** This capability is particularly advantageous for replicating physiological conditions and improving the functionality of tissue-engineered constructs, such as tubular structures in the human kidney, which require small-scale dimensions. Additionally, these microgels can be functionalized with biomolecules as outlined in **Scheme S3 (SI)**. This was exemplified by functionalizing microgels with an anesthetic drug procaine hydrochloride, enabling its sustained release over a period of 7 days (**Figure S17)**.

### 2.5 Outlook

Although a few studies on 3D and 4D (bio)printing with jammed microgels have been reported, they often aim to enhance shear properties to improve printability.^[37]^ Indeed, many of these approaches rely on the use of additional binder hydrogels to stabilize 3D structures.^[12,33,34,45]^ Moreover, the materials used in these studies often lack inherent bioactivity.^[34]^ In contrast, this study demonstrates 4D printed stimuli-responsive microgel-based constructs that do not require a secondary hydrogel as a binder, thereby leveraging microporous void structures for efficient nutrient transfer and vascularization, and rapid response to stimuli. Additionally, the microgels utilized in this study were inherently bioactive and can be further functionalized with therapeutic agents for sustained drug delivery, as well as with contrast-enhancing agents to improve magnetic resonance imaging visibility (MRI) as presented in **Scheme S4 and S18 (SI)**.

This dual-crosslinking strategy can be effectively extended to other protein biopolymers, including collagen, silk, keratin, decellularized extracellular matrix, and albumen, for carbohydrazide functionalization; and carbohydrate biopolymers such as cellulose, chitin, lactose, and starch for their oxidation potential, thereby broadening the scope of material synthesis. Consequently, this study opens multiple avenues for future research and development of interparticle crosslinkable and ion-responsive microgels in tissue engineering and regenerative medicine. For example, leveraging synthetic chemistry, this polymer can be engineered to facilitate controlled drug delivery, contingent on the gel’s degradation profile. Moreover, therapeutics encapsulated within these microgels can be precisely administered to target locations due to microgels’ injectability and ease of manipulation. As shown in **Scheme S3 and S4 and description in Section 2 (SI),** the -COOH and -NH_2_ coupling reactions can be used to functionalize the polymer, and microgels can be fabricated from that polymer. Using this technique, microgels can be functionalized with growth factors, small interfering RNAs (siRNAs), messenger RNAs (mRNAs), and microRNAs (miRNAs) for controlled spatiotemporal release,^[71,72]^ guiding protein expression, gene regulation, and stem cell differentiation into specific lineages. While our work highlights the potential of soft microgels of various sizes and shapes for creating dynamic constructs and programmable structures, further reinforcement with different materials can extend their applications to various tissues like muscle, tendon, cardiac, and bone. For instance, skeletal muscle tissue engineering can be accelerated with 4D bioprinted porous microgel-based constructs, which will not only maintain long-term cellular viability and facilitate early vascularization but also improve their functionality via ion-responsive mechanical actuation to train muscle during its maturation. Moreover, 4D printing with stimuli-responsive microgels offers exciting possibilities beyond tissue engineering, including soft robotics^[73]^, iontronic sensors^[62]^, biohybrid devices such as bioelectronics (wearable devices) with real-time environmental responsiveness^[74]^, and programmable materials for regenerative medicine and prosthetics.

## 3 Conclusion

In this study, we demonstrated a robust Schiff-base crosslinking mechanism, incorporating functionalization with hydrazide and methacrylate groups of protein biopolymers and oxidation of carbohydrates, to fabricate printable ion-responsive microgels. This approach obviates the need for temperature or enzymatic crosslinking, thereby preserving cellular viability and enabling bioprinting under physiological conditions. Interparticle photocrosslinking yielded stable 3D constructs with tunable mechanical and structural properties while retaining void spaces within these constructs. µR exhibited superior mechanical properties and packing density compared to µS. Consequently, µS-based constructs allowed the formation of thicker blood vessels in-ovo, while µR-based constructs supported the development of a denser vascular network. This distinction underscores the adaptability of these microgels for specific applications. Additionally, 3D bioprinting using µR was scalable, demonstrating properties such as shear thinning, self-healing, bioprintability, and high shape fidelity, where various bioprinting techniques, including EBB (in air), embedded and intra-embedded bioprinting, and AAB, were employed. These properties were coupled with inherent bioactivity and void space of microgels, making them promising candidates for tissue engineering, particularly in applications which require rapid vascularization. Furthermore, stimuli-responsive behavior of these microgels enabled 4D printing to develop programmable structures that underwent structural transformations into predicted shapes, such as coiling and folding, upon stimulation. Overall, this study establishes a foundation for bioactive, dual-crosslinked, self-supporting, bioprintable, and stimuli-responsive microgels, paving the way for the development of tissues and tissue models and devices such as biosensors or soft actuators for medical applications.

## 4 Materials and methods

### 4.1 Material synthesis and microgel fabrication

#### 4.1.1 Synthesis of OA

OA was synthesized by following a previously reported method.^[75]^ Briefly, 2 % (w/v) of sodium alginate (Sigma-Aldrich, St. Louis, MO, USA) solution was prepared by dissolving 4 g of sodium alginate in 200 mL DI water with the help of magnetic stirring at room temperature. Parallelly, 1.4 g of sodium periodate was dissolved in 14 mL DI water to prepare a 10 % (w/v) solution, which was then added dropwise to the sodium alginate solution. The mixed solution was kept under stirring in a dark condition. After 24 h, the reaction was terminated by adding 3 mL of ethylene glycol for 45 min. Subsequently, 2 g of sodium chloride (Sigma-Aldrich, St. Louis, MO, USA) was added to the final solution. After 30 min of stirring, the resultant solution was purified by using a dialysis membrane with a molecular weight cutoff (MWCO) of 6-8 kDa in DI water for 3 days, where water was replaced every day. Afterward, the retentate was frozen at -80 °C and lyophilized (FreeZone 18L Console Freeze Dry System, Labconco, Kansas City, MO, USA) for two days. The chemical structure of OA was verified using ^1^H NMR (AVANCE III 500 MHz, Bruker, MA) and attenuated total reflectance-Fourier transform infrared spectroscopy (ATR–FTIR, Nicolet 6700, Thermo Fisher Scientific, MA, USA) **(Scheme S2 and Figure S19 in SI)**.

#### 4.1.2 Synthesis of FG

GelMA was initially synthesized following a previously reported method.^[76]^ 3 g of the prepared GelMA was dissolved in 300 mL DI water. Parallelly, 450 mg 1-Ethyl-3-(3-dimethylaminopropyl)carbodiimide (EDC) (AK Scientific, Union City, CA, USA)was dissolved in 10 mL of water and slowly added to the GelMA solution. The mixture was then stirred continuously for 15 min. Then, 450 mg 1-Hydroxybenzotriazole (HOBt) (Sigma-Aldrich, St. Louis, MO, USA) dissolved in 10 mL dimethyl sulfoxide (DMSO) (Sigma-Aldrich, St. Louis, MO, USA) was added dropwise to the resultant solution. Subsequently, 2.2 g of carbohydrazide (Sigma-Aldrich, St. Louis, MO, USA) was added, and the pH was adjusted to 5.2 and the solution was stirred continuously overnight. The reaction mixture was dialyzed in 0.3 M NaCl solution, 25% ethanol, and pure water sequentially for varying periods (2 days in NaCl, 1 day in ethanol, 2 days in DI water) using a 6–8 kDa MWCO. The sponge form was then obtained via lyophilization. The chemical structure of FG was verified through ^1^H NMR and FTIR **(Scheme S1 and Figure S19 in SI)**.

#### 4.1.3 Schiff-base crosslinked bulk hydrogel

Bulk hydrogel was prepared by mixing FG and OA solutions. 5 and 10% FG solutions were made by dissolving FG in DI water at 50 °C, and 5 and 10% OA solutions were prepared by dissolving OA in DI water at room temperature. To obtain the crosslinked hydrogel, 5% or 10% FG was transferred into a 3D-printed cylindrical mold (4.35 mm in diameter, 2.35 mm in height) printed using an X-MAX printer (Qidi Tech, Zhejiang, China) with poly-lactic acid filament, and 5 or 10% OA was added into the same mold and mixed properly using a pipette 5 times. These hydrogel constructs were used for compression testing to optimize the concentration and ratio of OA and FG (**As described in Section 1.1 and shown in Figure S1 at SI).**

#### 4.1.4 Fabrication of µS

To generate spherical shaped microgels (µS), 10% FG and 5% OA were mixed in 4:1 ratio and injected into 75 mL mineral oil with 3% span 80 while being continuously stirred at 1,500 rpm with a magnetic stirrer for 30 min. Next, oil was placed into a 50 mL tube and thoroughly washed three times with DI water at a centrifugation speed of 2,904 ×g (Thermo Scientific, Sorvall Legend X1R), and the supernatant was removed, which yielded µS in bulk amounts.

#### 4.1.5 Fabrication of µR

To fabricate randomly shaped microgels (µR), a solution containing 10% (w/v) FG and 5% (w/v) OA was prepared at a volumetric ratio of 4:1 and allowed to undergo crosslinking for 30 min. Next, the hydrogel was then transferred into a blender with DI water and crushed for 2 min. The microgel solution was transferred to a 50 mL centrifuge tube and subjected to centrifugation at 2,904 ×g for 5 min in a centrifuge to pellet µR. The pelleted microgels were washed three times with DI water by repeating the centrifugation step at 2,904 ×g for 5 min, with the supernatant carefully removed after each wash.

### 4.2 Physicochemical characterization

#### 4.2.1 Microgel size

To quantify the size of microgels, a Mastersizer 3000 (Malvern Panalyticals, Worcester, UK) was used to determine the volume-weighted mean particle size. Both µR and µS, prepared separately three times to get triplets, using the above-mentioned methods, and the microgel size distribution was measured accordingly.

#### 4.2.2 Pore size, and porosity

Initially, microgels were fabricated without fluorescein isothiocyanate-dextran (FITC) to ensure it could later be used as an external tracer to measure the void space between microgels. These microgels were dispersed in a 5 mg/mL FITC-dextran (2 MDa) solution and mixed on a shaker for 5 min, followed by centrifugation at 19,632 ×g. The microgels were then loaded into a 3D-printed cylindrical mold (6 mm in diameter, 1.2 mm in height) printed using the X-MAX printer, which was attached to a thin glass slide with Sylgard 184. After transferring microgels, the mold was covered with glass slide to prevent dehydration. Images were captured in the z-direction using a LSM 880 Zeiss confocal microscope (Thornwood, NY, USA) with a 10x objective lens to visualize FITC-dextran within the interstitial pores of microgels. FIJI ImageJ (National Institute of Health, Bethesda, MD) was used to analyze porosity and pore characteristics. The image stacks were converted to 8-bit format and threshold based on the fluorescence signal intensity. Particle analysis was performed, with each identified particle corresponding to a two-dimensional pore. The analysis generated data for void fraction (porosity), pore count, and pore cross-sectional area, with these values averaged across all 40 images.

#### 4.2.3 Rheological characterization

Rheological characterization of µS and µR was performed using a rheometer (MCR 302, Anton Paar, Austria). All measurements were taken three times using a 25 mm parallel-plate geometry, kept at 22 °C with the help of a Peltier system. A flow sweep test was performed to evaluate the shear thinning behavior of both microgel types, where the shear rate varied from 0.1-100 s^-1^. Amplitude sweep was performed to determine viscoelastic properties at a constant angular frequency of 1 rad s^-1^ while shear strain varied from 0.001 to 1000 %. To investigate the self-healing behavior of µS and µR, a recovery sweep test was performed, where a change in viscosity was evaluated with low and high shear rates of 0.1 s^-1^ for 60 s and 100 s^-1^ for 10 s, respectively, and this was repeated two times.

#### 4.2.4 Mechanical characterization

To understand the mechanical properties of uncrosslinked and crosslinked microgels, a compressive test was performed. Firstly, microgels with LAP (5 mg/mL (w/v)) were transferred to a cylindrical mold (7.57 mm in diameter, 3.65 mm in height) printed using the X-MAX 3D printer and crosslinked to form into a 3D construct and transfer to the rheometer. Next, the plate was moved to ensure full contact with the construct. Compression was applied at a rate of 0.001 mm s^−1^. For cyclic testing, 20% strain was applied and released repeatedly (three times) to evaluate the construct’s behavior under repeated loading (**Figures S2A-B**). For failure stress and strain analysis after crosslinking, 80% strain was applied to assess the constructs’ fracture limits. The compressive modulus was calculated from the slope of the initial linear region of the stress versus strain curve for each sample.

#### 4.2.5 Fourier-transform infrared (FT-IR)

Microgels were prepared using emulsion and blending techniques as described before. These microgels were taken into two parts, one part was crosslinked, and another part was left uncrosslinked. Next, microgels were freeze-dried using a freeze dryer. FT-IR spectroscopy was performed using a diamond attenuated total reflectance (ATR) instrument. The spectra were collected within a wavenumber range of 4,000–400 cm⁻¹, with a resolution of 6 cm⁻¹, to analyze the chemical composition and functional groups present in microgels.

#### 4.2.6 Nuclear magnetic resonance (NMR)

Gelatin, GelMA, FG and OA powder were dissolved in deuterated water (D_2_O) to make solutions with a concentration of 10 mg mL^−1^ solutions, which were then transferred to an NMR tube. Next, proton NMR spectra were acquired using a Bruker AVIII-HD-500 (Billerica, MA, USA) spectrometer equipped with a Bruker Ascend magnet operating at 500 MHz.

#### 4.2.7 3D Printing characterization

##### 4.2.7.1 Hanging filament test

Firstly, µR microgels were prepared, and LAP solution (5 mg/mL) was added to these microgels, subsequently centrifuged at 19,632 rpm for 5 min. The resultant solution was used for 3D bioprinting purposes. µR were utilized as a bioink and evaluated for hanging filament length and assessed for their extrusion fidelity using 23, 25 and 27 G nozzles. The bioink was transferred into a 3 ml syringe barrel and then extruded using controlled pneumatic pressure, which was maintained at 40-180 kPa with the help of an INKREDIBLE+ bioprinter (CELLINK, Sweden). For maximum hanging filament length, a Nikon D810 camera with Nikon 105 Micro lens was used to record videos of filament extrusion. The bioink formed a filament while coming out from the nozzle and the maximum filament length was measured before the filament broke. After recording the video, the frame at which the filament detached from the nozzle was captured, and the filament length was measured using ImageJ.

##### 4.2.7.2 Filament fusion test

3D Bioprinting was performed to conduct the filament fusion test, printability, and spreading analysis according to the literature.^[39,77]^ Briefly, square pores of different sizes from 1 × 1 to 5 × 5 (mm × mm) were printed on a glass slide using µR. µR was extruded through 23, 25 and 27 G nozzles at bioprinting speed of 150 mm s^−1^ and extrusion pressure of 40-180 kPa. After printing square pores, images were taken using a camera and ImageJ was used to calculate pore area and perimeter. Material spreading (*S*) was determined using Equation (1), providing a measure of the extent of bioink spreading post extrusion. A percent diffusion value of ’0’ indicated no spread, meaning that the actual area matched the theoretical area. Here, *At* and *Aa* represented the theoretical and actual pore areas, respectively. Additionally, the printability (*Pr*) of bioinks was assessed using Equation (2), with a value of ’1’ signifying optimal printability

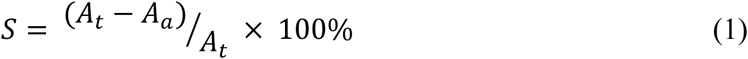

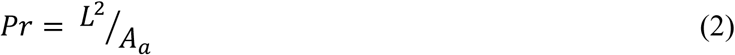

In Equation (2), ’*L*’ represents the perimeter of the printed pores.

##### 4.2.7.3 Filament collapse test

The filament collapse test was conducted by printing filaments across a linear array of pillars positioned at distances of 1, 2, 3, 4, 5, 6, 7, 8, and 9 mm. After printing, photographs were taken, and ImageJ was used to calculate the collapse area factor (*CF*), defined as the percentage of deflected area relative to the theoretical area, using Equation (3).

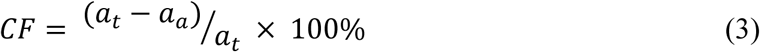

Where *a_a_* and *a_t_* represented the actual and theoretical areas, respectively. *a_a_* was considered zero for collapsed filaments, resulting in *CF* being 100%.

#### 4.2.8 Degradation

Percent degradation (*D* (%)) of µR and µS constructs was measured by submersing them in PBS at 37 °C and changing the PBS solution every day. The constructs were taken out at various time points, namely Days 1, 2, 3, 7, 14, and 28 and gently rinsed with Milli-Q water, and dried completely. *D* (%) was calculated using the equation below:

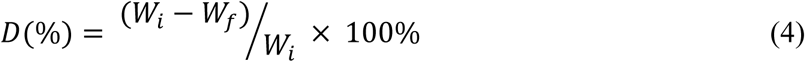

Where *W_f_*and *W_i_* represented the dry weight after Days 1, 2, 7, 14, and 28 and the initial weight of the constructs, respectively.

### 4.3 3D Bioprinting of microgels

#### 4.3.1 3D Bioprinting organ models

To fabricate organ models, such as ear, pancreas and nose, µR based bioink was used. Initially, a mesh file (downloaded from https://sketchfab.com/) of the nose and ear was sliced to generate G-codes using a slicer, and then these organ models were bioprinted (INKREDIBLE+ bioprinter) using µR mixed with RhB dye solution (2 mg/mL) with a 23G nozzle at constant printing speed of 600 mm/min and extrusion pressure of 80 kPa.

#### 4.3.2 Microgels as a support bath

To characterize microgels as support material, 1% (w/v) sacrificial XG was used for printing (using the INKREDIBLE+ bioprinter) into the support bath of µR and µS. First, the sacrificial XG was loaded into a barrel and a 23G needle was used to study the correlation between filament diameter and printing speed by incrementally increasing the print speed from 100 to 300 mm/s at 60, 70, 80 and 90 kPa extrusion pressures. Images were taken using a tabletop EVOS microscope (Thermo Fisher Scientific, US). These images were analyzed using ImageJ. Moreover, this embedded printing of XG in µR was used to print complex shapes such as spiral and branched structure, which was followed by removal of XG and perfusion with RhB dye.

#### 4.3.3 Intra-embedded printing

A pancreas model was fabricated using the intra-embedded printing technique^[56]^, where the material served dually as both a bioink and support bath. Briefly, the pancreas model was digitally processed and sliced to generate the corresponding G-code for embedded printing. The model was subsequently printed with µR using the INKREDIBLE+ bioprinter within a 1.5% (w/v) XG support bath, utilizing a constant printing speed of 600 mm/min and an extrusion pressure of 100 kPa to ensure precise filament deposition and structural fidelity during fabrication. Following this, intra-embedded printing of 1% (w/v) XG mixed with different color oil dyes (Winton Oil Color, London, UK) was performed within the µR-based pancreas model, using custom-written G-code, to achieve a nested vascular structure.

#### 4.3.4 Cell culture and viability test

Cell experiments were conducted in a Biosafety Level-2 safety cabinet (LabGard® ES Class II, Type A2, Plymouth, MN). GFP^+^MDA-MD-231 breast cancer cells were provided by Dr. Danny Welch, from the University of Kansas (Kansas City, KS) (passage 30–35) were cultured with µR, µS, and bulk hydrogels to test the material toxicity and cell attachment. The culture medium consisted of Dulbecco’s Modified Eagle Medium (DMEM) (Thermo Fisher Scientific, USA) enriched with 10% fetal bovine serum (FBS) (Life Technologies, Grand Island, NY, USA), 1% penicillin-streptomycin (PS, Life Technologies), and 1 mM glutamine (Life Technologies, Carlsbad, CA, USA). For cell viability, cells were cultured for 1, 3, and 7 days and viability were calculated using LIVE/DEAD staining. Ethidium homodimer-1 (EthD-1, 4 μM) dye (Invitrogen, CA, USA) was used to stain dead cells, while GFP fluorescence was used for staining the viable cells on Days 1, 3, and 7. Before staining, cells were rinsed three times with Dulbecco’s phosphate-buffered saline (DPBS) (Corning, USA). The samples were observed under an Axio Observer microscope (Zeiss), and images were acquired after 1h of incubation (*n* = 3 per day). ImageJ was used to calculate cell viability. Further, NHLF (Lonza, Houston, Texas, USA) and HDF (ATCC: The Global Bioresource Center, USA) were also cultured in µR and µS, where DMEM was supplemented with 10% Plenty (Plenty Bio, Thousand Oaks, CA, USA) to understand the biocompatibility of these microgels with different types of cells.

#### 4.3.5 Aspiration-assisted bioprinting (AAB) characterization

A custom-designed AAB platform was used to bioprint spheroids within prepared µS and µR baths, as reported previously,^[57,78]^ to further investigate the impact of microgel shapes on bioprinting accuracy, precision and cellular migration. In this regard, prepared µS and RM were transferred into the designed 3D printed cubic devices (7.35 mm in length, 7.35 mm in width, and 1.2 mm in height), serving as supporting baths, with the device boundaries used as reference points for AAB. The prepared GFP^+^MDA-MB-231/HDF spheroids were bioprinted inside µS and µR at predefined target positions with known distances in between. After bioprinting, the actual distance between bioprinted spheroids and the reference point was measured under the EVOS microscope, and a total of 5 spheroids were bioprinted and subsequently analyzed for each microgel bath type. The accuracy of bioprinted

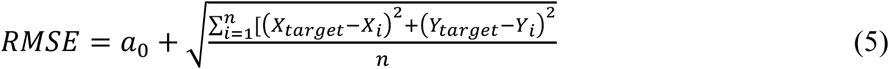

The root mean square error (RMSE) was used to quantify the accuracy of AAB process. Here, *X_target_* and *Y_target_* denote the target position coordinates, while *X_i_* and *Y_i_* represent the coordinates of the bioprinted spheroids along the x-and y-axes, respectively. The sample size was indicated by *n*. Precision was assessed as the square root of the standard deviation of the measured positions. A lower value for both accuracy and precision metrics indicates higher printing accuracy and precision.

#### 4.3.6 AAB for spheroid proliferation

Firstly, 3D printed molds with a mold diameter of 6 mm and height of 1 mm were fabricated, using an X-max 3D Printer, which was subsequently sterilized by soaking them in 70% ethanol and subjecting them to UV light for 1 h. Parallelly, multicellular spheroids were formed by seeding a cell suspension of GFP^+^MDA MB-231 and HDF at a 2:1 ratio in a 96-well plate at a density of ∼6,000 cells per spheroid, followed by 24-h culture. AAB was used to bioprint spheroids into the 3D-printed molds (6 mm in diameter and 1.2 mm in height) filled with µS and µR. For BH constructs, FG and OA were combined with spheroids in the molds which was further crosslinked with 405 nm light. The constructs were cultured in DMEM supplemented with 10% Plenty, 1 mM glutamine, and 1% penicillin-streptomycin at 37 °C with daily medium changes. After culturing for 1, and 7 days, the constructs were fixed in 4% paraformaldehyde (PFA) followed by PBS washing. Tissue clearing solution, prepared through a previously reported method^[79]^, was used to clear the constructs, and z-stacks of cell outgrowth were obtained using a Zeiss LSM 880 confocal microscope. Maximum intensity projection images were generated using ImageJ to quantify spheroid area.

#### 4.3.7 CAM assay

The impact of microporous architectures within the synthesized microgels on angiogenesis was assessed utilizing the chick embryo CAM assay. In brief, fertilized chick eggs, kindly provided by Poultry Education and Research Center at Penn State (University Park, PA, USA), were incubated in a horizontal position at 37.8 °C in a 60% humidified controlled atmosphere using a universal egg hatching incubator (Peatend, China) with automatic rotation, and the start of incubation was designated as embryonic Day 0. Following 7 days of incubation, a circular window was created at the air chamber between eggshell and CAM under sterile conditions, and then the prepared samples were gently implanted on top of CAM of the chick embryo. Subsequently, the windows of eggs were sealed with sterilized transparent parafilm tape, and the eggs were returned to the incubator and further incubated without the rotation setting. On Day 14, the transparent parafilm tape was removed, and the implanted samples were harvested and imaged with a camera (Pixel 8, Google, USA), followed by fixation with 4% PFA. Then, embryos were euthanized by adding 4% PFA (5 mL) into eggs. All the operational and inoculation processes were illustrated in **Figure 3A**. To quantitatively analyze the total number of neo-vessels, images of the inoculation area on the CAM, where the implanted constructs were located, were processed using ImageJ and Angio tool (**Figure S8**).

#### 4.3.8 Histological Assessment and Immunohistochemistry

CAM assay samples were fixed within 4% PFA at room temperature (RT) for 24 h, followed by dehydration in 15% sucrose solution for 5-6 h and then 30% sucrose solution for 12 h. Samples were embedded in O.C.T. cryomatrix (Epredia, Kalamazoo, MI, USA) and sectioned at 10-µm thickness using a Leica CM1950 cryostat (Leica Biosystems, Deer Park, IL, USA) at -20 °C. The cryosections were used for histological and immunofluorescence staining. For H&E staining, sections were processed using a Leica Autostainer ST5010 XL, mounted, and visualized with the Zeiss Axiozoom V16 inverted microscope. For immunofluorescence staining, cryosections were thawed at RT for 15 min from -30 °C, washed in Milli-Q water and PBS (15 min each), permeabilized with 0.2% Triton X-100 for 10 min, and blocked with 5% Bovine Serum Albumin (BSA) (Sigma-Aldrich, St. Louis, MO) for 1 h at RT. Vessel detection was performed by incubating sections with Lycopersicon esculentum (Tomato) Lectin (Invitrogen, 1:500 in 1% BSA) for 1 h, followed by PBS washing. Counterstaining was done with DAPI for 10 min, followed by additional PBS wash. Sections were mounted with ProLong™ Gold Antifade Reagent containing DAPI and imaged using the Zeiss Axio Observer microscope.

### 4.4 4D Printing

#### 4.4.1 Shrinkage characterization

Microgels were first produced as described before. Next, µS were kept in DI water and images were acquired using the EVOS microscope before and after the addition of 1X PBS. These images were then used to calculate the change in diameter of the microgels using ImageJ. Further µS, µR and BH constructs were prepared by casting them into a cubic shape mold (7.35 mm in length, 7.35 mm in width, and 1.2 mm height) and crosslinked using 405 nm as described earlier. To study the kinetics of shrinkage, 1X PBS was added, and videos were recorded by time. This was repeated three times per sample. After 280 s, PBS was removed, and these constructs were kept in DI water to study the recovery, and the time of the complete recovery was recorded. The same study was conducted for µR with 0.5M, 5M sucrose, DMSO, 137 mM NaCl, 6M NaCl, 1X PBS at 4 °C, 37 °C, pH 1, and pH 14 and 10X PBS (at room temperature and pH 7.4) to study the effect of other stimuli, such as temperature, concentration, and pH, on the shrinkage kinetics. Further, 4M LiCl, NaCl, KCl, CaCl_2_ and MgCl_2_ (Sigma-Aldrich, St. Louis, MO, USA) solutions were used to treat µR constructs to further understand the shrinkage mechanism.

#### 4.4.2 Fabrication of GelMA microgel ink

GelMA microgels were fabricated using a blending technique. Briefly, a 15% (w/v) GelMA solution was prepared in deionized (DI) water, and 5 mg/mL LAP was added as the photoinitiator. The GelMA solution was then crosslinked under 405 nm light and blended for 2 min using a blender (Magic-bullet, Los Angeles, CA, USA). The resulting solution was centrifuged at 4,182 ×g for 5 min to pellet the microgels. The microgels underwent three wash cycles with deionized (DI) water to thoroughly remove any uncrosslinked GelMA. Subsequently, the prepared microgels were combined with a 20% (w/v) GelMA hydrogel solution at a 1:1 volumetric ratio. The mixture was centrifuged at 19,632 ×g for 5 min, and the supernatant was removed. The resultant GelMA microgel ink was used for 4D printing.

#### 4.4.3 Multilayered 4D printing

For 4D printing, a multilayer strategy was used, where one layer was printed using GelMA microgel ink prepared as described in Section 4.4.2 and the second layer was printed using µR, and these two layers were crosslinked using 405 nm light for 2 min. Further, these printed constructs were transferred into a 1X PBS bath for 10 min, and images were taken. This strategy was used to print coiling filaments, grippers, and folding sheets.

#### 4.4.4 4D Physics-based modeling

##### 4.4.4.1 Model fitting

Shrinking and non-shrinking microgels were incompressible (i.e. no change of volume under deformation) and highly deformable in a non-linear elastic fashion. In this regard, a 2-parameter Mooney-Rivlin material model was utilized that accounted for the non-linear elasticity to fit the experimental compression tests. To incorporate the Mooney–Rivlin material model, engineering strain (*ε*) was converted to stretch ratio (*λ* = *ε* + 1). The experimental compressive stress (*σ*) was used, which is a component of the first Piola-Kirchhoff stress tensor along the direction of sample deformation. The formulation for the first Piola-Kirchhoff stress response was derived for the case of uniaxial extension or compression of the incompressible Mooney-Rivlin material, capturing the material’s behavior under such loading condition as follows ^[80]^:

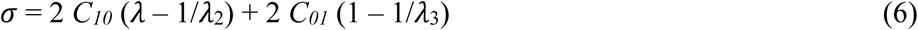

Here, *C_10_* and *C_01_* were the microgels parameters for a Mooney–Rivlin material. The curve-fitting toolbox from MATLAB (Natick, MA) was used for curve fitting with a minimization of the least squares of the error (R^2^ = 0.99). C_10_ and C_01_ were determined to be 0 and 1839.5 Pa for the shrinking microgels and 321.35 and 5068.9 Pa for the non-shrinking microgels, respectively. In the small strain regime, the shear (*μ*) and elastic modulus (*E*) for incompressible microgels were calculated from these material constants as:

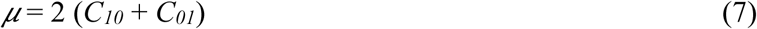

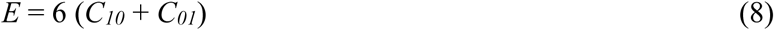

##### 4.4.4.2 Calibration of the shrinkage coefficient

A thin square-shaped film (7.3 cm in width) of the shape-changing material was submerged into 1X PBS and allowed to shrink to its maximum extent (6.3 cm average observed width after shrinkage). In general, commercial software does not include packages for microgels swelling or shrinkage but mostly they do include thermal expansion and shrinkage. In this regard, the heat transfer node can be coupled with solid mechanics node to provide the constitution of microgels shrinkage and thermal stresses. An analogy can be used for change in temperature with the change in swelling media, thermal stresses with swelling stresses, thermal expansion strains with swelling strains.^[81]^ In this work, COMSOL Multiphysics version 0.2 (COMSOL Multiphysics, USA) with solid mechanics, heat transfer and thermal expansion Multiphysics coupling was utilized. A 2D square with a side of 7.3 cm was designed, and the temperature was decreased by 1 K (Kelvin) to calibrate a thermal coefficient that resulted in a shrinkage to a width of 6.3 cm. A value of 0.1 K⁻¹ was determined for the thermal coefficient. All simulation studies were conducted in a steady state and a uniform distribution of temperature throughout the entire domain was assumed. A density of 1,000 kg/m³, thermal conductivity of 1,000 W/(m·K), and a heat capacity at constant pressure of 1 J/(kg·K) were used in simulations. The elastic properties of shrinking microgels were obtained from the Mooney-Rivlin fit.

##### 4.4.4.3 Finite Element Modeling of 4D materials

For simulation of different shapes, computer-aided design (CAD) files mimicking the printed bilayer structures were created. Material properties were assigned according to different microgel layers, either shrinkable or non-shrinkable. The temperature was then decreased to observe the resulting change in shape. These studies were conducted using a stationary solver and a numerical technique known as the method of continuation. Instead of solving a single problem by decreasing the temperature by 1 K, a small fraction of the temperature (0.1 K) was decreased and a stationary solution was obtained. The solution obtained was used as the initial condition for the subsequent step, in which the temperature was further decreased by another fraction. This method was continued until the total temperature change of 1 K was achieved. In this manner, the final deformed shapes of the printed microgels were obtained.

## Statistical Analysis

All data were presented as mean ± standard deviation (SD). Data were plotted using OriginPro 2021b (Northampton, MA). Statistical differences were determined using one-way analysis of variance (ANOVA) followed by Tukey’s post hoc test. Results fulfilling the null hypothesis at *p* < 0.05 were considered statistically significant (*), while at *p* < 0.01 (**), p < 0.001 (***) and p < 0.0001 (****) as highly significant and ‘ns’ represents not significant.

## Supporting information

Description of Additional Supplementary Files

## Acknowledgements

This work was supported by the National Institute of Biomedical Imaging and Bioengineering (NIBIB) Award R01EB034566 (I.T.O.) and the National Institute of Dental and Craniofacial Research (NIDCR) Award R01DE028614 (I.T.O.). We extend our gratitude to Dr. Jian Yang (Penn State) for providing access to material characterization resources, Plenty Bio (Thousand Oaks, CA, USA) for generously providing Plenty through the Beta Program, Dr. Thomas Neuberger for assistance with MRI data acquisition, and Annie Smith for her help with histology. We also thank Mr. Scott L. Kephart from the Poultry Education and Research Center at Penn State University, who kindly provided fertilized chicken eggs for the CAM assay.

## Notes

### Competing Interest Statement

I.T.O. has an equity stake in Biolife4D and is a member of the scientific advisory board for Biolife4D and Healshape. The remaining authors declare no competing interests.

