## Supplementary information for "Interparticle Crosslinked Ion-responsive Microgels for 3D and 4D (Bio)printing Applications"

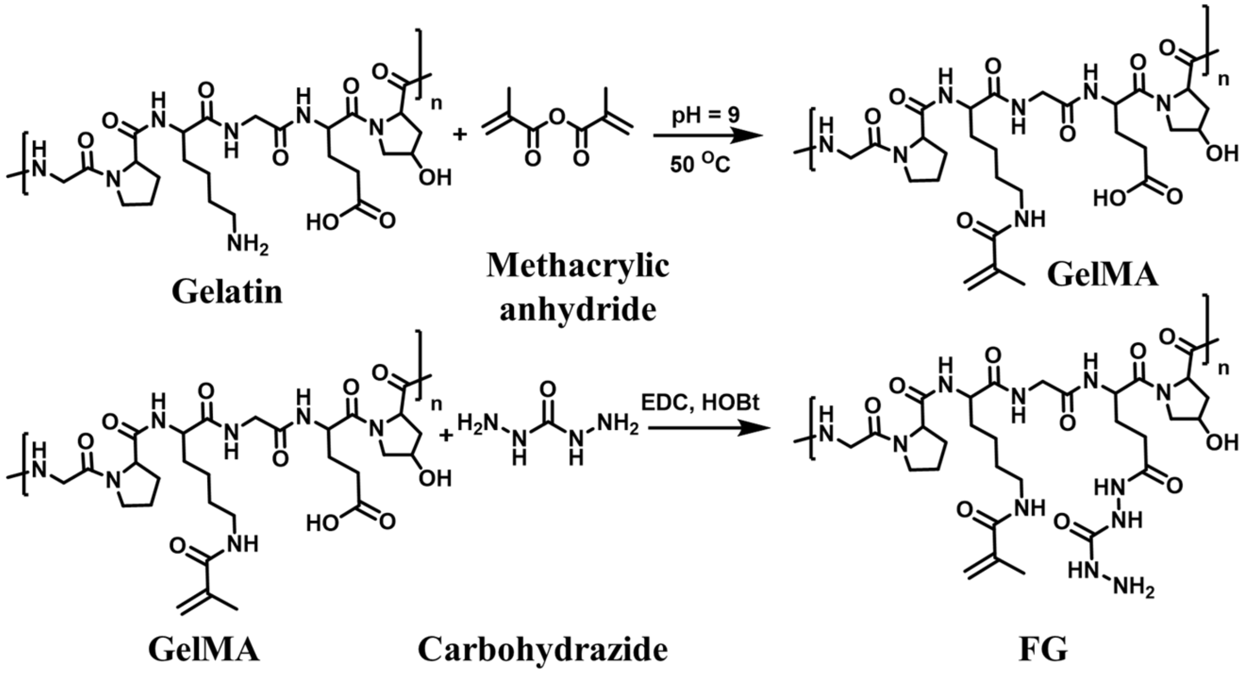


**Scheme S1.** Synthetic scheme of FG.

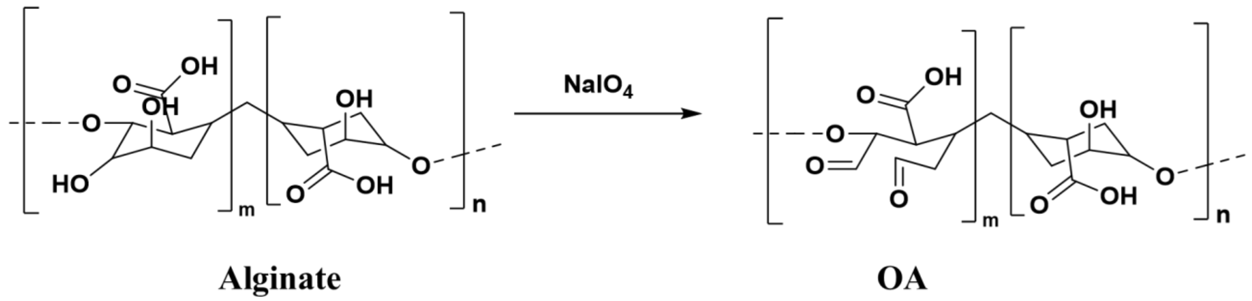


**Scheme S2.** Synthetic scheme of OA

#### 1.1. Compression test of the Schiff-base crosslinked hydrogel

##### 1.1.1 Optimization of concentration of FG and OA

To perform the compression test, cylindrical molds (4.35 mm in diameter, 2.35 mm in height), 3D printed using the X-MAX 3D printer, were used to shape the bulk hydrogels, which were prepared by mixing FG and OA at different concentrations. Hydrogel formulations were prepared by mixing solutions of FG (5 and 10% concentrations) and OA (5 and 10% concentrations). Equal volumes of each solution were combined at a 1:1 ratio, resulting in four different crosslinked hydrogel formulations. The compression experiments were conducted using the Anton Paar rheometer equipped with a 1 kg load cell. Each sample was placed on the rheometer, and the plate was moved to contact the sample, setting the zero position. The samples were then compressed at a rate of 0.005 mm $s^{-1}$ until reaching 70% strain. The compressive modulus was calculated from the linear region of the stress versus strain curve (**Figs. S1 A1 & A2**). The compression tests revealed that the hydrogel prepared with 5% OA and 10% FG exhibited superior mechanical properties with a compressive modulus of 12.06 ± 0.11 kPa, which was significantly higher than other composition, as illustrated in **Figs. S1 A1-A2**.

##### 1.1.2 Optimization of the mixing ratio of FG and OA

Initially, the individual concentration of FG and OA was optimized. Following the concentration optimization, the mixing ratio of FG and OA was optimized by evaluating four sample types with the following OA:FG ratios: (1:9), (1:4), (3:7), and (2:3). The compression test results indicated that the 1:4 ratio provided the highest mechanical properties, with a significantly higher compressive modulus of 67.24 ± 8.40 kPa, as shown in **Figs. S1 B1-B2.**

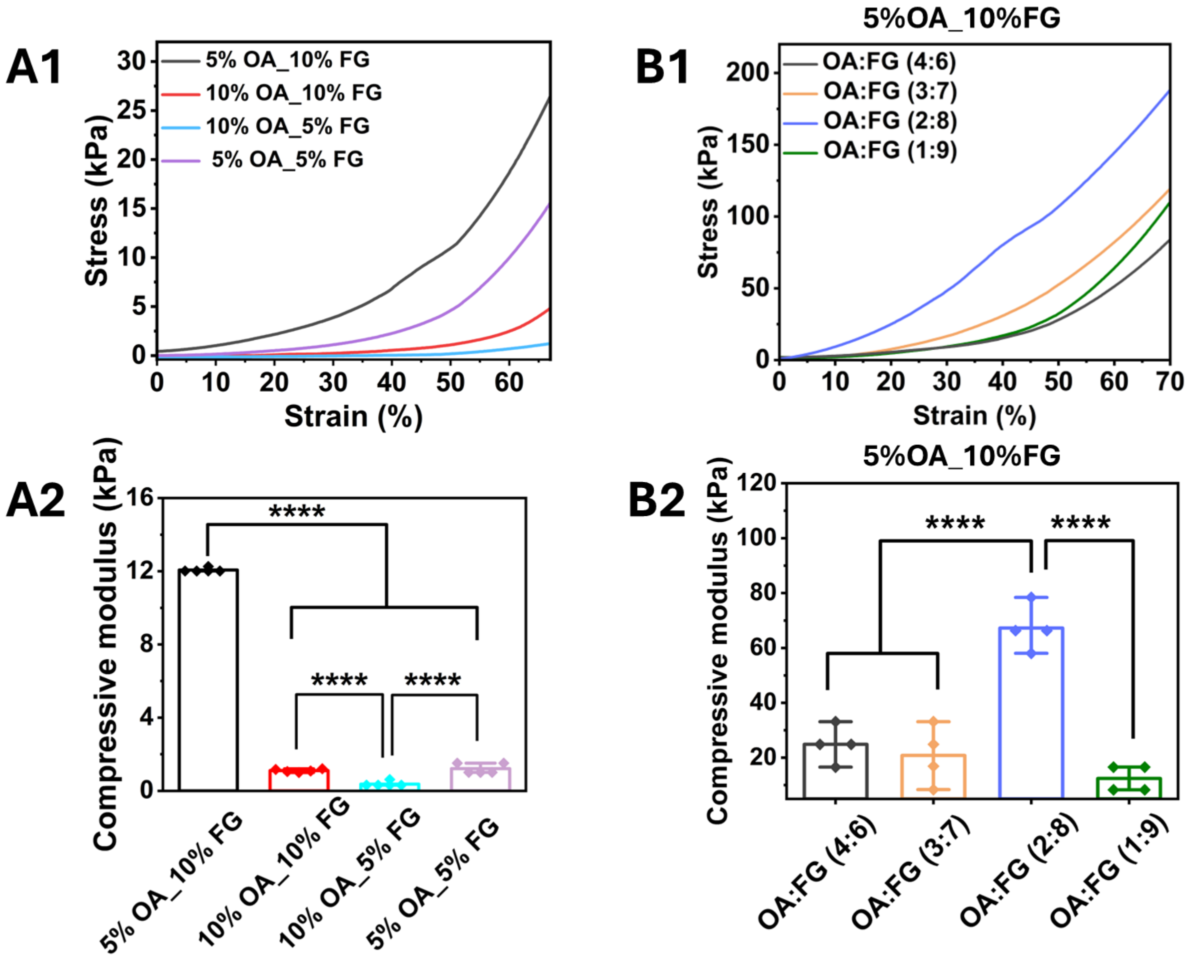


**Figure S1.** Evaluation of the effect of different concentrations and ratios of OA and FG on mechanical properties of fabricated microgels: (A1-B1) the representative stress-stain curves of different samples with different OA and FG concentrations and ratios, and (A2-B2) the corresponding compressive modulus (mean ± SD, $n \geq 4$, $^{****}$ $p \leq 0.0001$).

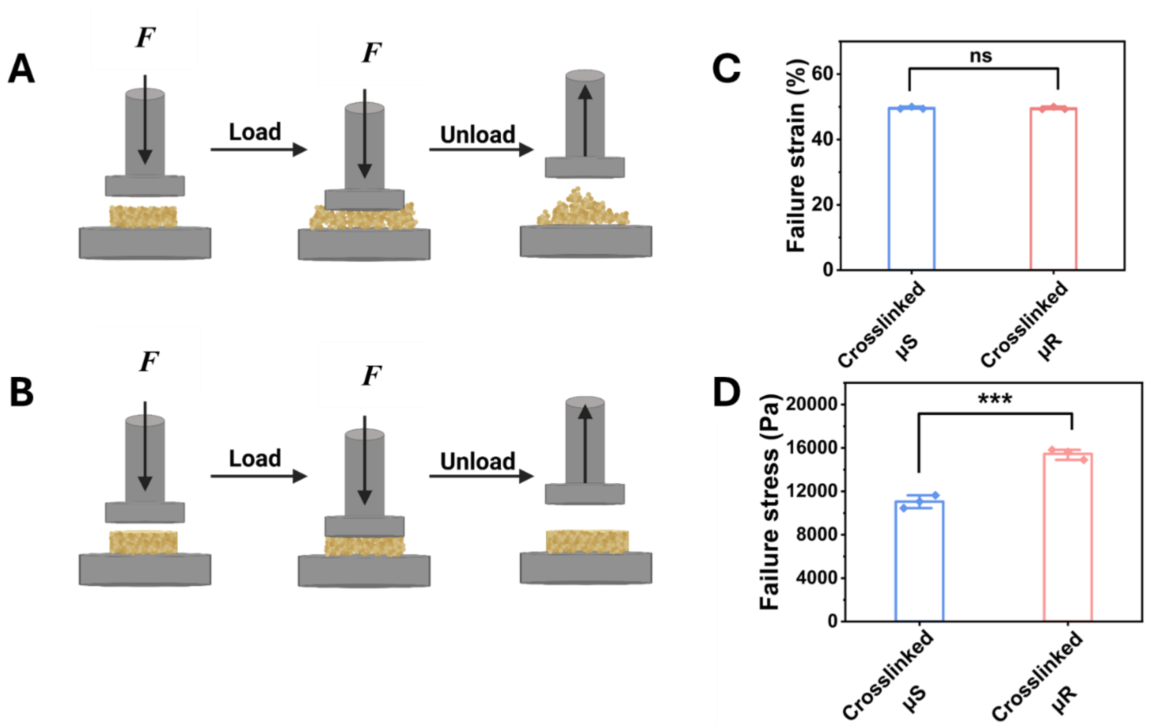


**Figure S2**. Characterization of mechanical properties of uncrosslinked and crosslinked μR and μS constructs (7.57 mm in diameter, 3.65 mm in height). Schematic representation of cyclic compression process and corresponding sample deformation with and without applied forces (*F*) in (A) uncrosslinked, and (B) crosslinked microgels. Stress and strain measurement of both μS and μR constructs under compression up to 80% or until failure where (C) failure strain at the breaking point and corresponding (D) failure stress (mean ± SD, $n$=4, $^{***}p \leq 0.001$, ns indicates non-significant).

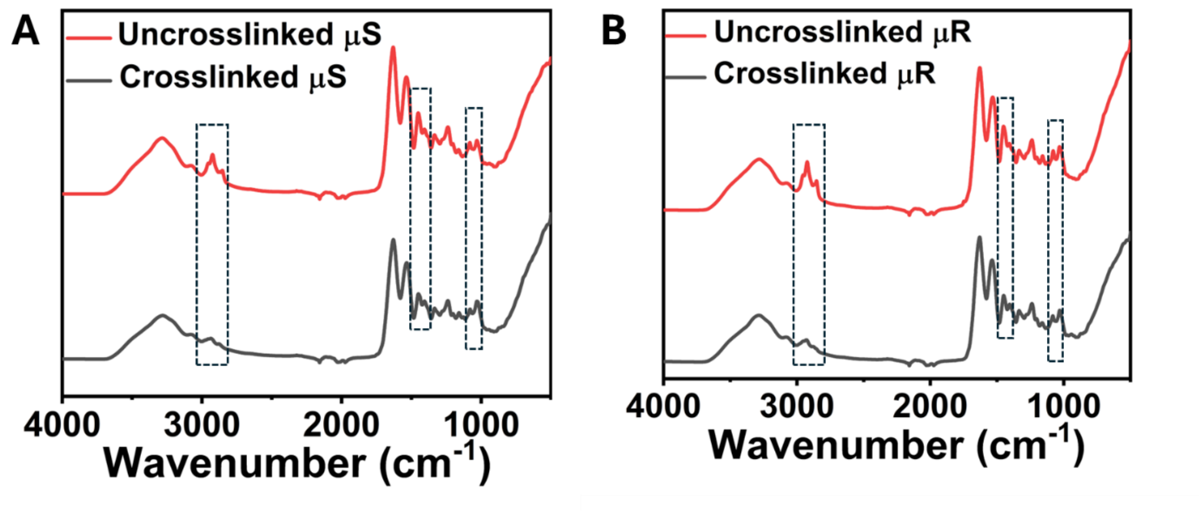


**Figure S3.** The FTIR spectra of uncrosslinked and crosslinked (A) μS and (B) μR.

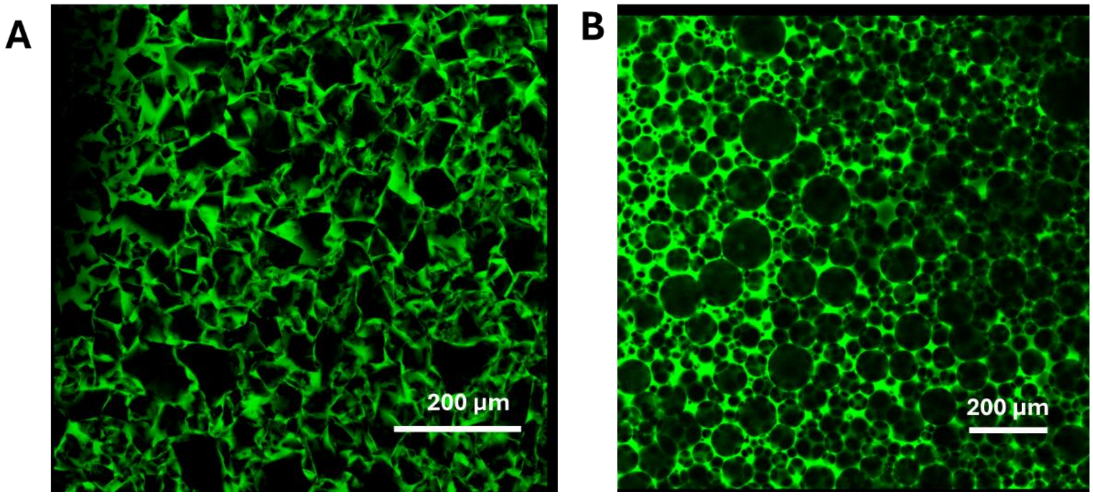


**Figure S4.** Fluorescent images of porous structures within (A) µR and (B) µS constructs (green: FITC-Dextran, presented porous structure and area, black area presented microgels).

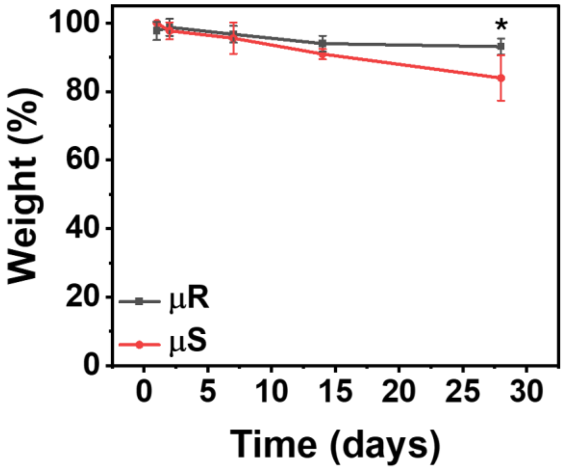


**Figure S5**. Evaluation of degradation of μR- and μS- based scaffolds samples in PBS (mean ± SD, *n*= 4, $^{*}p \leq 0.05$).

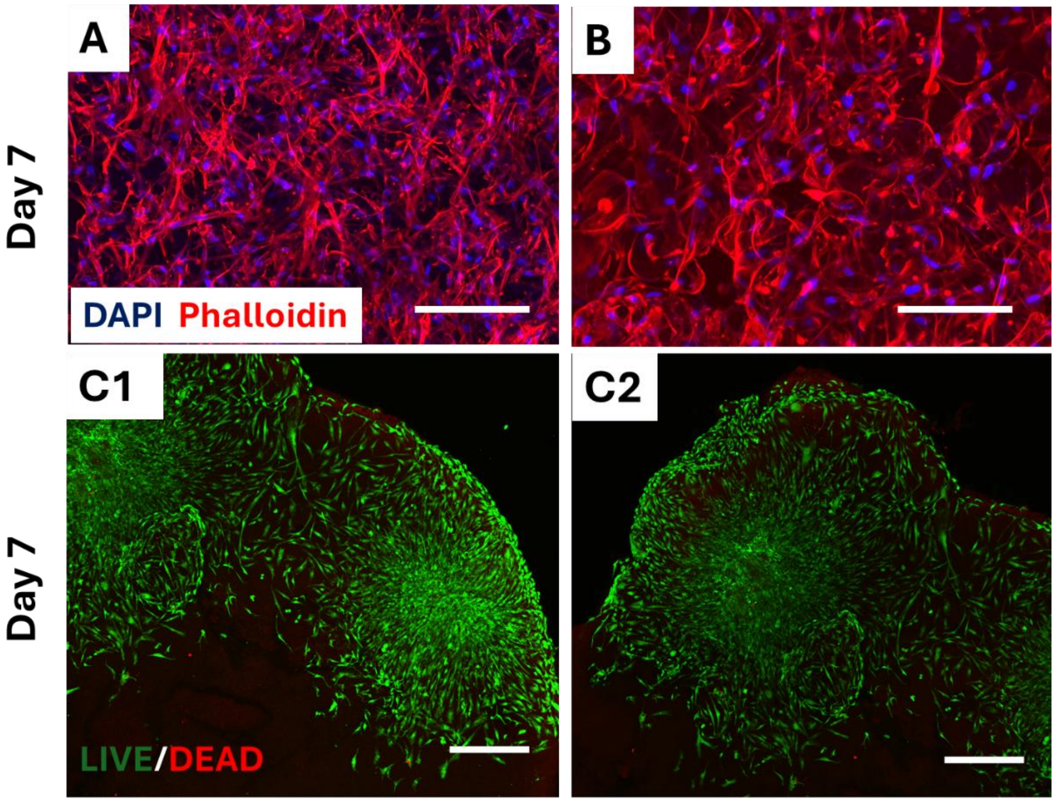


**Figure S6.** Representative DAPI/Phalloidin fluroscence images of proliferation of NHLF cells grown in (A) μR-based and (B) μS-based construct at Day 7 (scale bar – 500 μm). (C1-C2) LIVE/DEAD fluorescence images of HDF spheroids spreading in μR-based construct at Day 7 (scale bar – 500 μm).

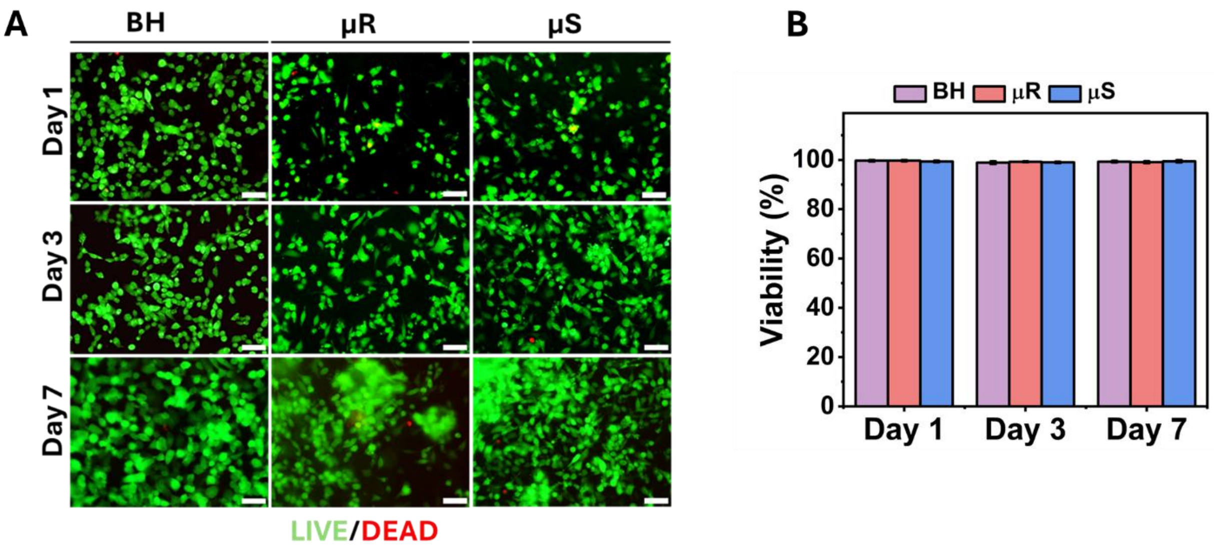


**Figure S7.** Adhesion, proliferation and viability of GFP$^+$MDA-MB-231 cells seeded on BH, μR, and μS constructs after 1, 3, and 7 days: (A) Representative LIVE/DEAD fluorescence images and (B) the corresponding cell viability (mean ± SD, *n* = 3).

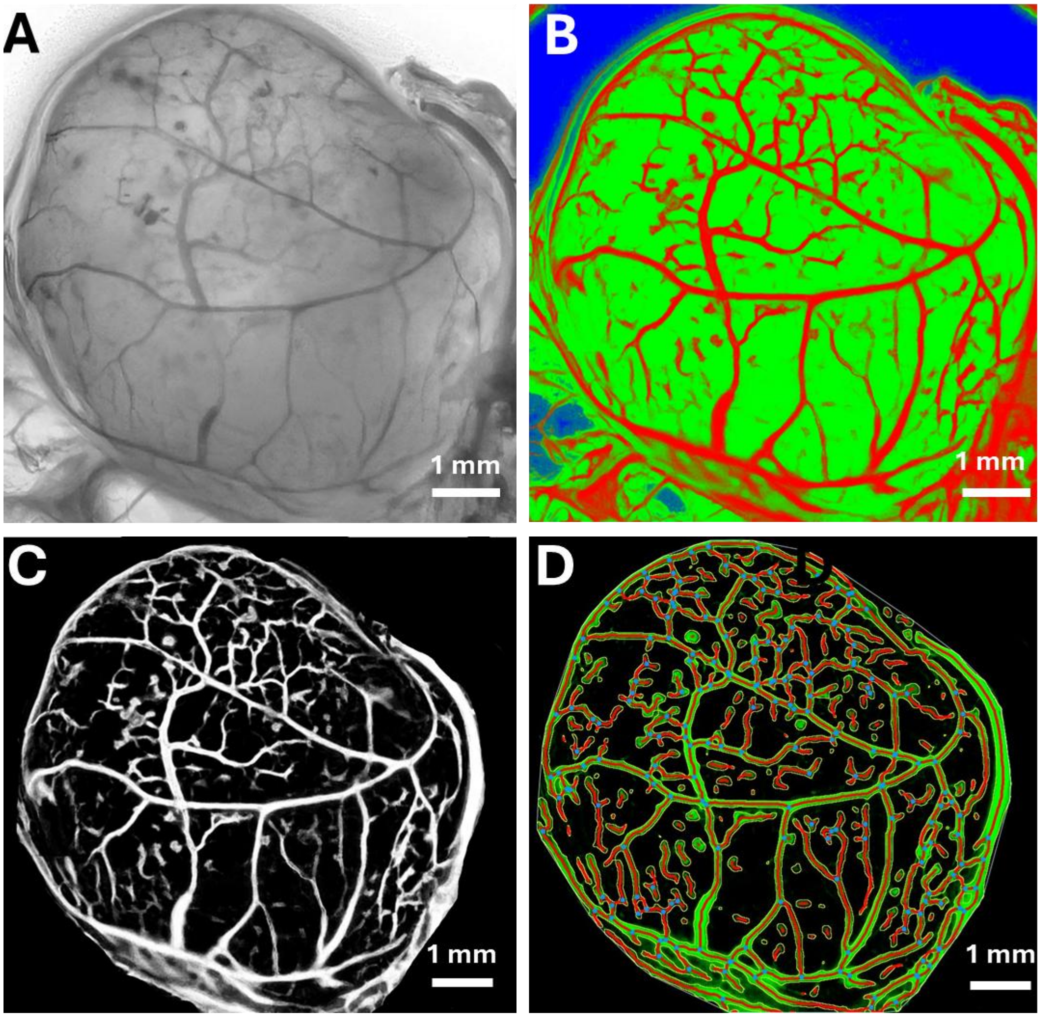


**Figure S8.** Representative analysis of harvested CAM samples: (A) a grayscale image of harvested μR constructs. (B) A colored image, where vessels were shown in red, and the construct was shown in green. (C) A binary image of the same construct. (D) An image generated using Angio tool, which gave information related to the total vessel length, vessel density, number of junctions and junction density.

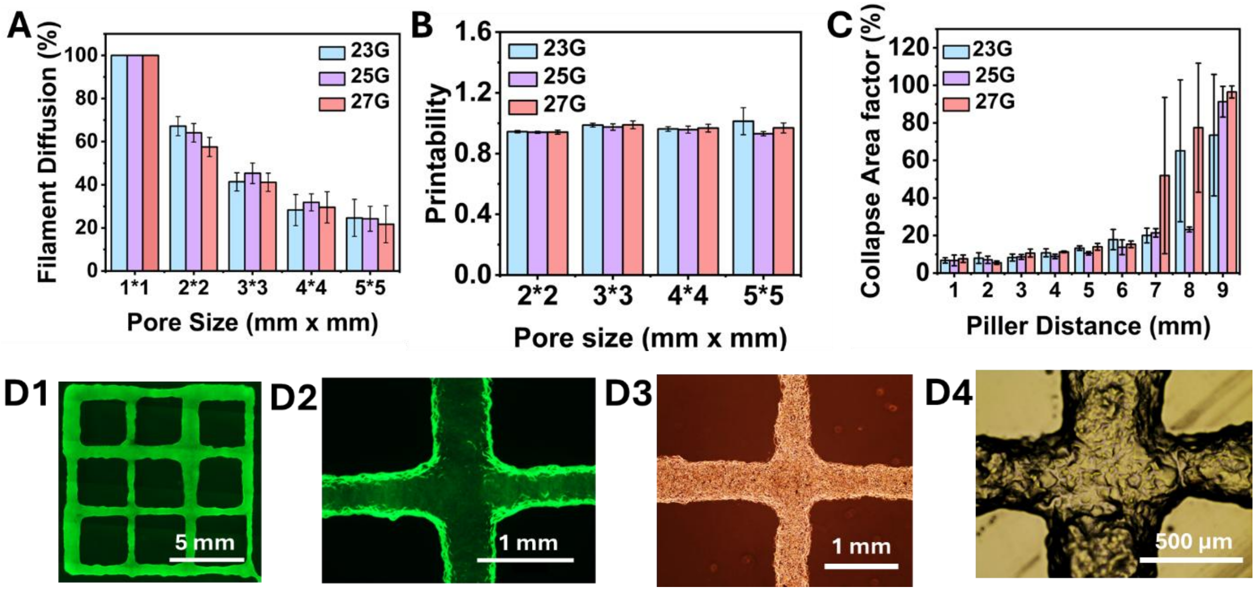


**Figure S9.** Printability and shape fidelity of μR-based ink with different nozzles: (A) printability, (B) filament fusion test and (C) collapse measurements. (D1-D4) Representative fluorescent images, where $GFP^{+}$MDA-MB-231 cells were mixed with μR, and bright-field images of printed porous constructs (mean ± SD, $n \geq 3$).

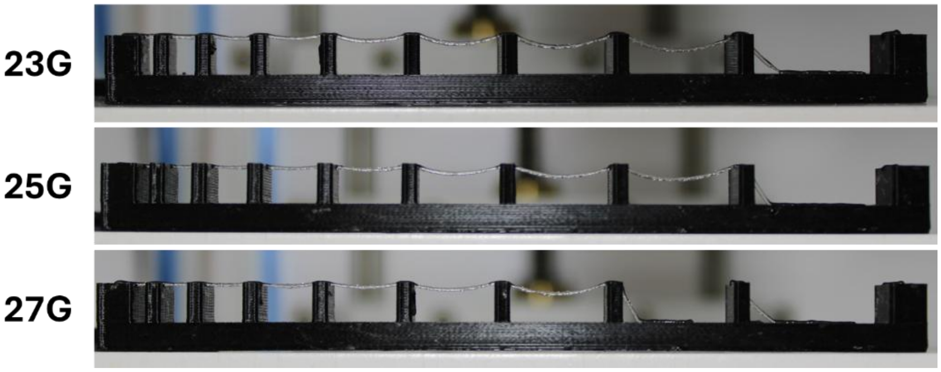


**Figure S10**. Photographs taken during the collapse test for filaments made of μR-based ink and printed with different sized nozzles.

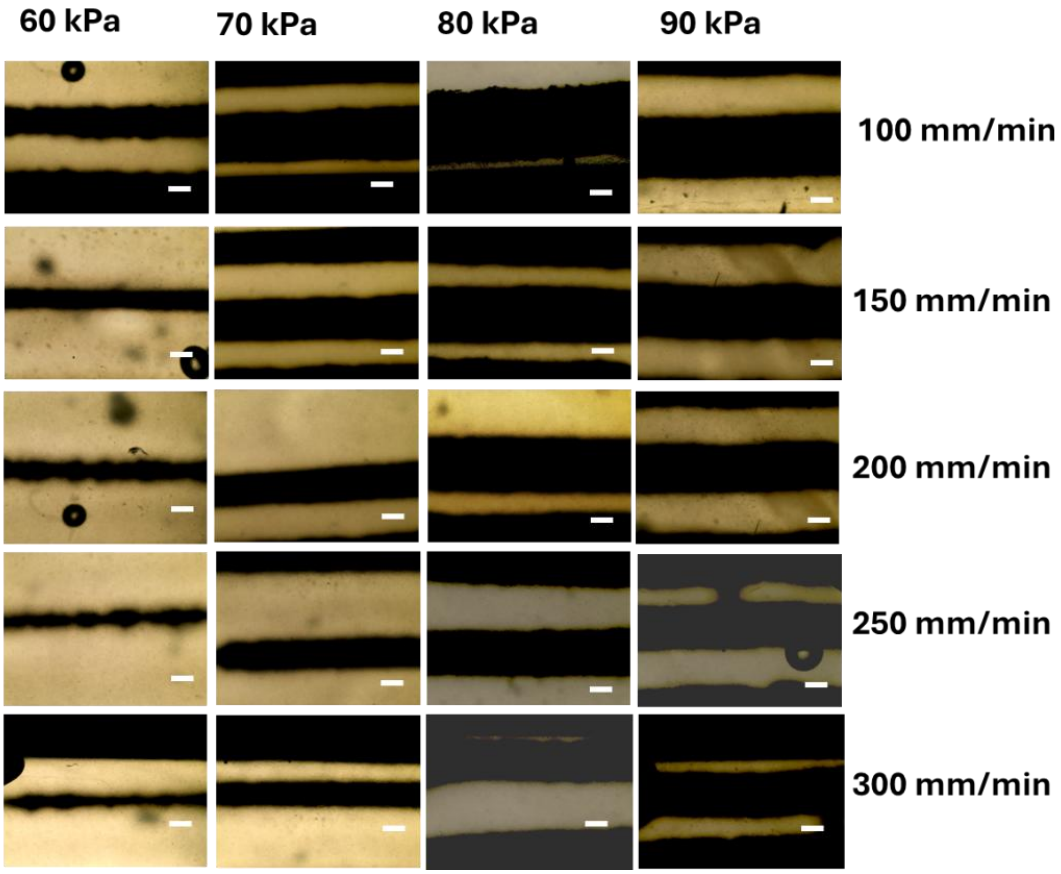


**Figure S11.** Evaluation of printability during embedded bioprinting using μR as a support bath material: photographs of representative printed filaments under different printing speeds and pressures (scale bar: 500 μm).

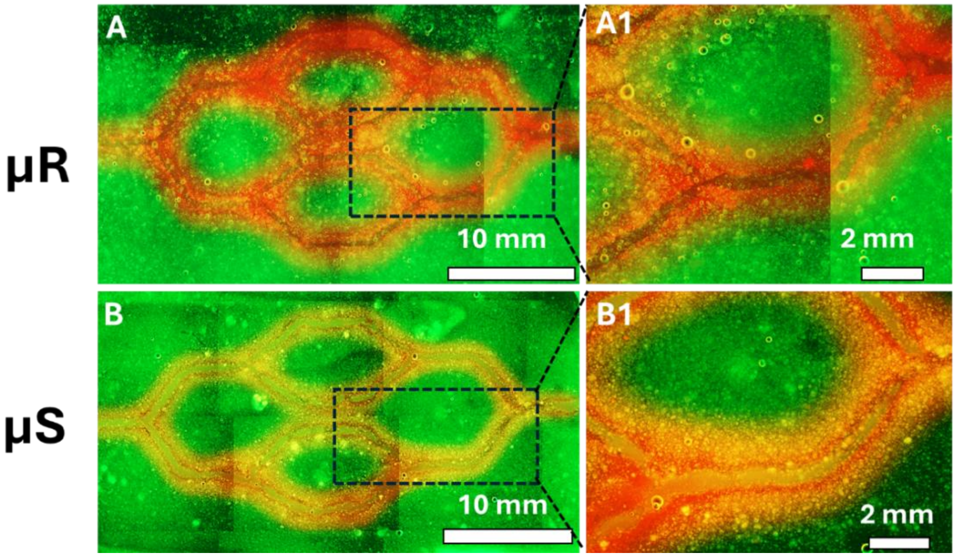


**Figure S12.** Construction of vasculature-like channels using 1% XG within GFP+ MDA-MB-231 cell loaded (A) μR and (B) μS as support baths for embedded bioprinting (green: MD-MB-231, red: red fluorescence represents rhodamine dye perfused after removing XG sacrificial ink).

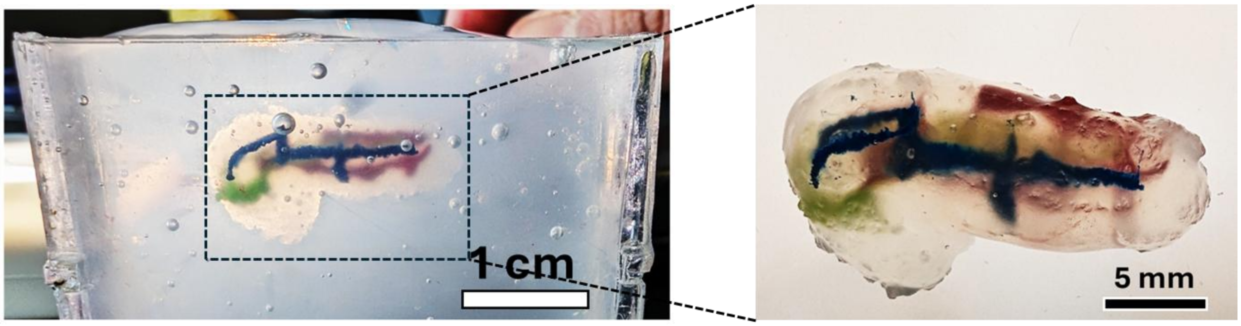


**Figure S13.** Photos of the printed complex pancreas model using the μR ink along with a vascular network printed using intraembedded bioprinting, in a support bath of 1.5% XG.

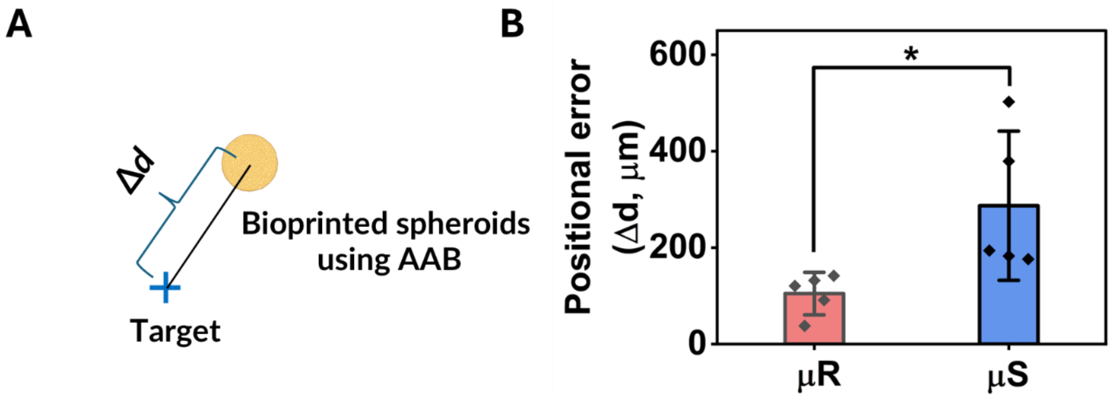


**Figure S14.** Schematic diagram and accuracy characterization of AAB of HDF/GFP$^{+}$MDA-BM-231 spheroids within μR and μS microgels: (A) a schematic for quantitative assessment of the bioprinting accuracy (Δ*d*: positional error) and (B) the corresponding bioprinting accuracy (mean ± SD, $n = 5$, $^{*}p \leq 0.05$).

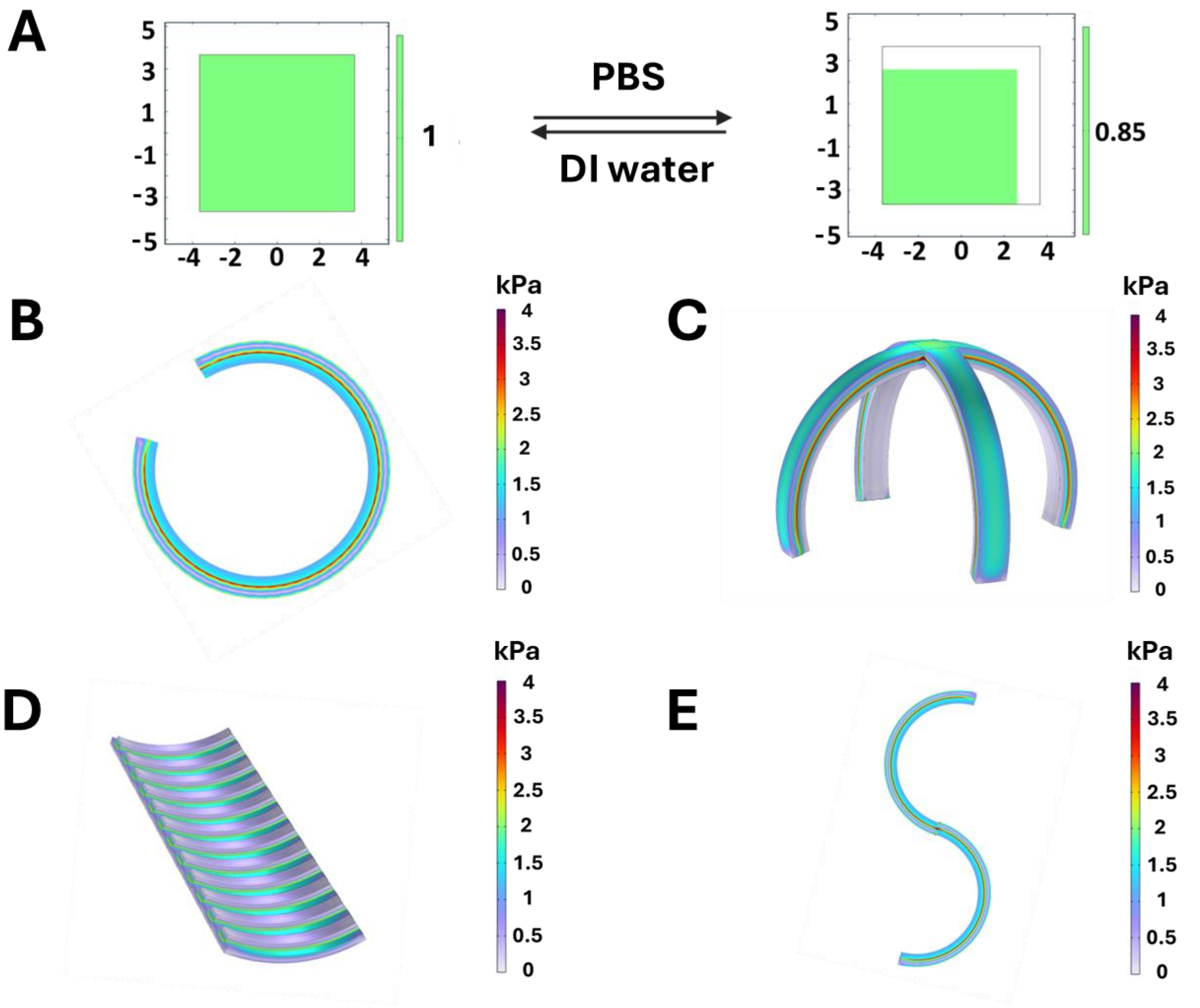


**Figure S15.** Simulated stress profiles of 3D printed multilayer structures and 4D transformation effect after stimuli response: (A) calibration of the simulation model, (B) a bi-layered straight line segment coiling into a circular shape, (C) a cross-shaped bi-layered structure transforming into a gripper, (D) a flat structure having horizontal lines exhibiting sheet folding, and (E) the transformation of a straight filament into an S-shape upon stimulation.

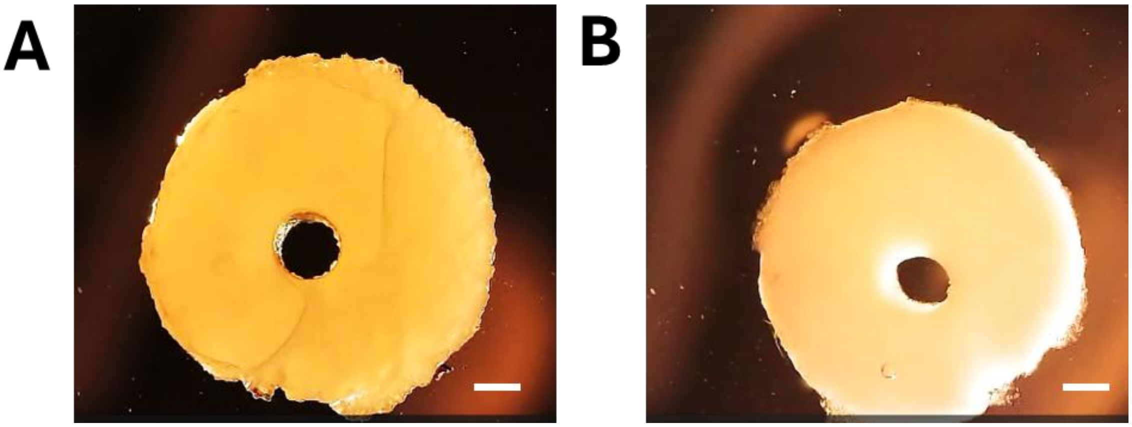


**Figure S16.** Shrinkage of a channel in μR-based constructs: (A) Before the addition of PBS, (B) after 10 min of PBS addition (scale bar – 7.5 mm).

### 2. FG Functionalization

#### 2.1. Functionalization with an anesthetic drug

To functionalize FG with a drug, 1 g GelMA was dissolved in 100 mL DI water. Further, 150 mg 1-Ethyl-3-(3-dimethylaminopropyl)carbodiimide (EDC) was dissolved in 3 mL water and added to the GelMA solution slowly and the solution was stirred for 15 min. Then, 150 mg hydroxybenzotriazole (HOBt) was dissolved in 3 mL DMSO and added dropwise to the resultant solution. Subsequently, 100 mg of procaine hydrochloride was added to the solution, and the pH was adjusted to 5.25. The solution was stirred for 6 h. Following this, 800 mg of carbohydrazide was added, the pH was readjusted to 5.25, and the mixture was stirred overnight. The reaction mixture was dialyzed to purify in 0.3 M NaCl solution for 2 days, 25% ethanol for 1 day, and pure water for 2 days using a 6–8 kDa MWCO. The drug loading efficiency was calculated using a nano-drop (Thermoscientific, Wilmington, DE, USA) after dialysis, yielding 30% based on Equation S1. The pure product was then obtained through freeze drying.

$$DE\ (\%) = {}^{DL}/_{TD}\, x100 \qquad (1)$$

Where Drug Loading Efficiency (*DE*) refers to the percentage of the drug that was successfully loaded into a solution after dialysis. *DL* represents the weight of the drug presented in the solution after dialysis, and *TD* denotes the total weight of the drug initially added to the solution before the reaction.

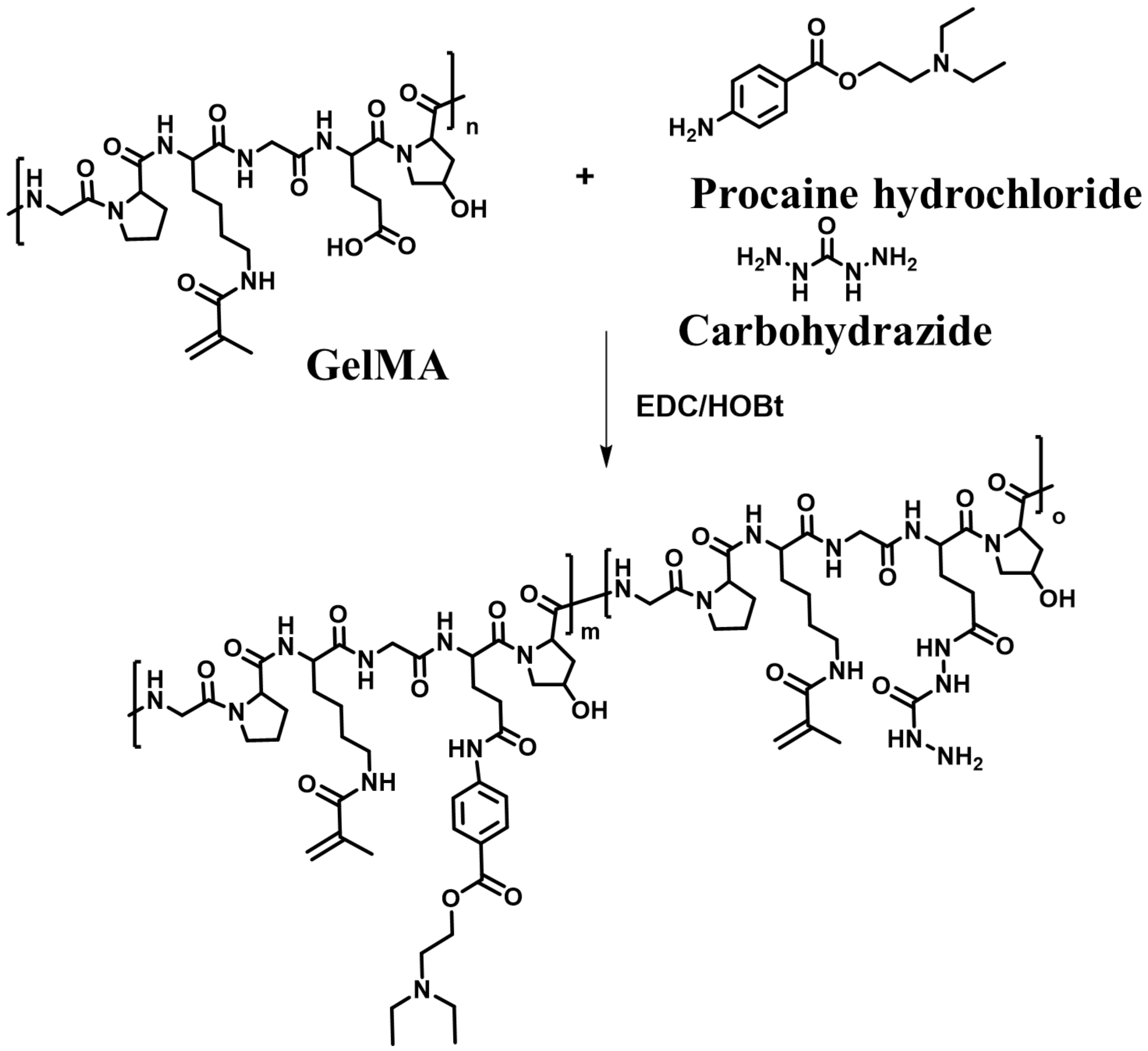


**Scheme S3.** Synthetic scheme of FG with procaine hydrochloride functionalization.

#### 2.2. Anesthetic drug release

To study the release of procaine hydrochloride, μR constructs were prepared and kept in PBS at 37 °C under shaking conditions. Samples were collected at Day 1, 3, 5 and 7. Nano-drop was used to measure the percent drug release **(Figure S17).**

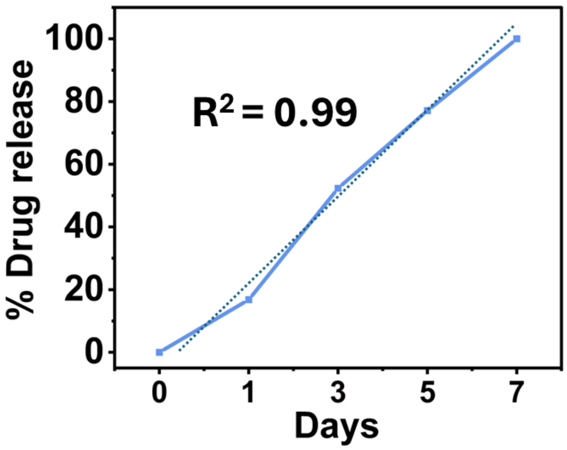


**Figure S17**. Procaine hydrochloride drug release profile from μR scaffold over a period of 7 days, where dash line shows the sustain release profile of the drug with $R^2$=0.99.

**2.3. Functionalization of the FG with a Magnetic resonance imaging (MRI) contrast agent**

To functionalize FG with an MRI contrast agent, 1 g FG was first dissolved in 100 mL DI water. Further, 100 mg diethylenetriaminepentaacetic acid gadolinium(III) dihydrogen salt hydrate was taken separately in 10 ml DI water and 150 mg EDC was dissolved in 3 mL water and added to Diethylenetriaminepentaacetic acid gadolinium(III) dihydrogen salt hydrate solution slowly and the solution was stirred for 15 min. Then, 150 mg of HOBt was dissolved in 3 mL of DMSO and added dropwise to the reaction mixture, which was stirred for 1 h. Subsequently, this solution was combined with the FG solution, the pH was adjusted to 5.25, and the mixture was left to react overnight under stirring. Following this, 800 mg of carbohydrazide was added, the pH was readjusted to 5.25, and the solution was stirred overnight. The resulting reaction mixture was purified by dialysis against 0.3 M NaCl solution for 2 days, 25% ethanol for 1 day, and pure water for 2 days using a 6–8 kDa MWCO. The pure product in sponge form was achieved after three days of freeze-drying.

Further, 100 mg of functionalized FG was dissolved in 1 ml DI water along with 5 mg LAP. This solution was transferred to a mold (7.57 mm in diameter, 3.65 mm in height) printed using X-MAX 3D printer and crosslinked with 405 nm light to generate cylindrical disk. This disk was visualized in MRI and given a sharp difference as compared to the control (pristine FG). (**Fig. S18**)

This process can be further optimized to get Schiff-base crosslinked microgels and implanted into body, which can be easily monitored using MRI.

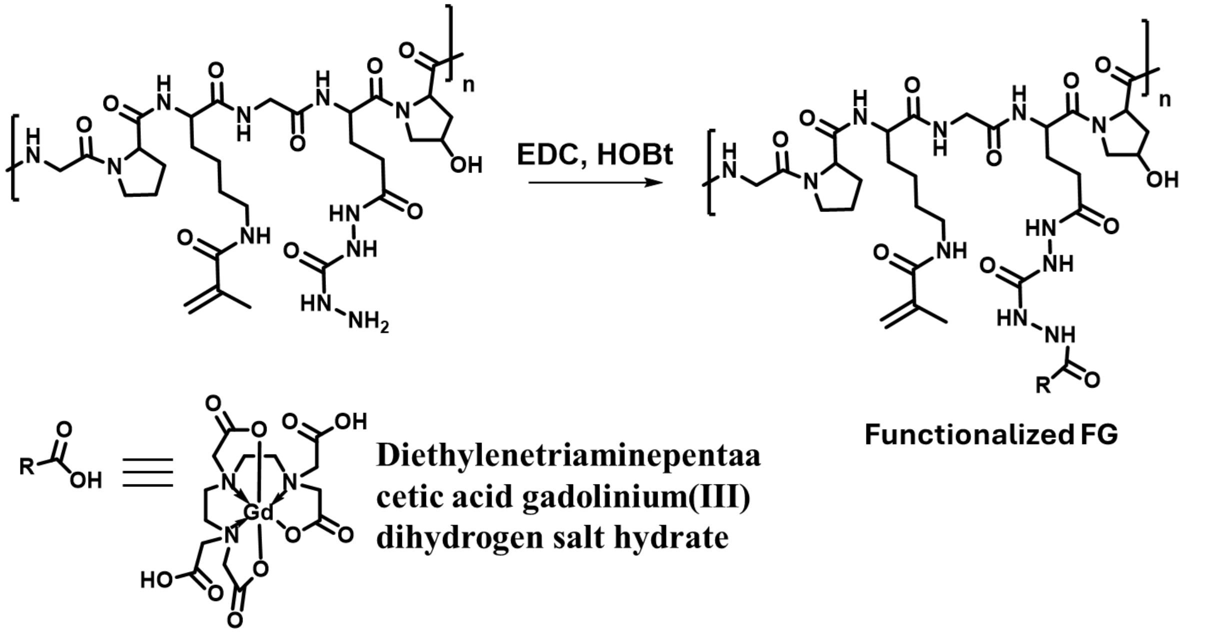


**Scheme S4**. Synthetic scheme of FG functionalization with diethylenetriaminepentaacetic acid gadolinium(III) dihydrogen salt hydrate.

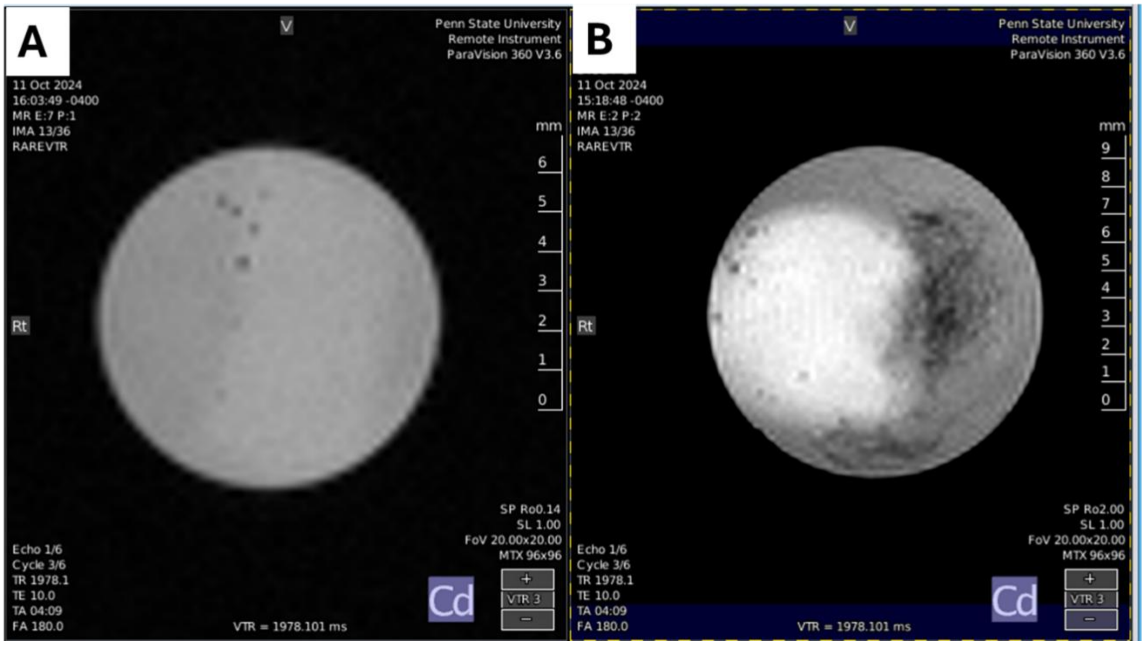


**Figure S18** MRI imaging of functionalized (A) pristine FG and (B) FG functionalized with MRI contrast agent.

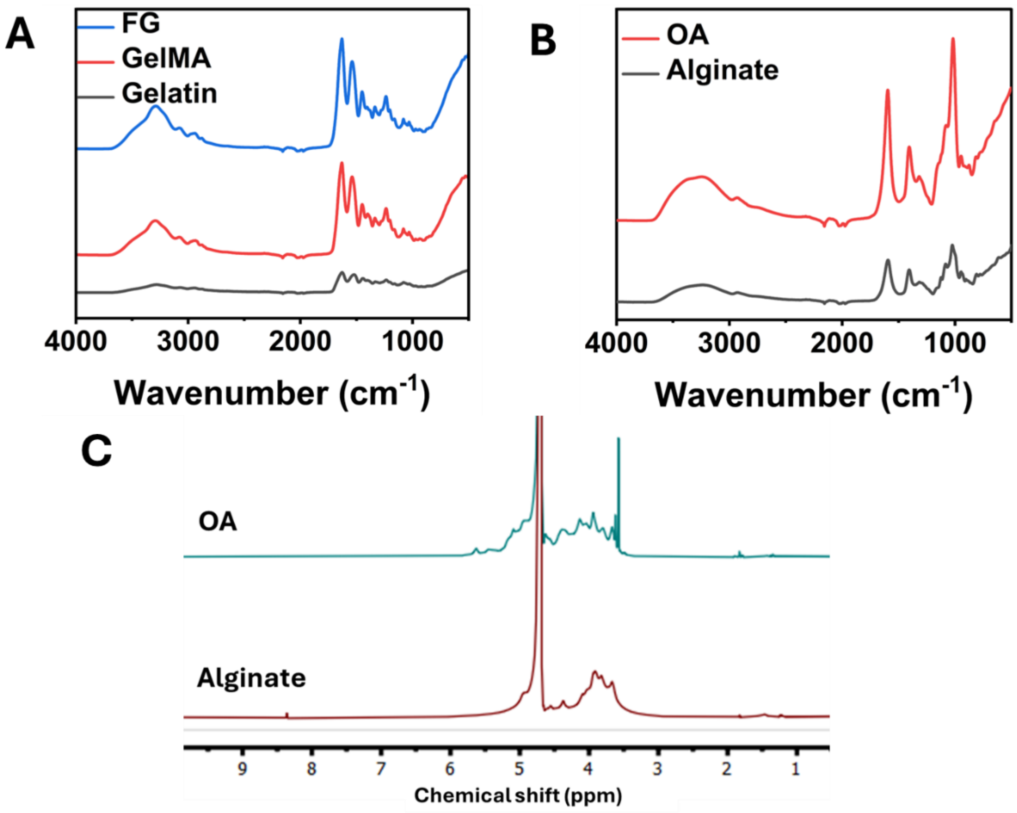


**Figure S19.** FTIR and NMR of synthesized polymers: (A) FTIR of FG, GelMA and gelatin, (B) FTIR of OA and alginate, (C) $^{1}$H-NMR of OA and alginate.
