## Supplementary material for "Interparticle Crosslinked Ion-responsive Microgels for 3D and 4D (Bio)printing Applications": Description of Additional Supplementary Files

**File Name:** Supp Video S1
**Description:** Reconstructed z-stack images for qualitative analysis of porosity and packing density of µR construct.

**File Name:** Supp Video S2
**Description:** Reconstructed z-stack images for qualitative analysis of porosity and packing density of µS construct.

**File Name:** Supp Video S3
**Description:** Fabrication of centimeter-scale ear using 3D bioprinting of µR followed by interparticle crosslinking.

**File Name:** Supp Video S4
**Description:** Shrinkage of µS after the addition of 1X PBS.

**File Name:** Supp Video S5
**Description:** Shrinkage of BH, µS and µR constructs after treating with 1X PBS.

**File Name:** Supp Video S6
**Description:** Shrinkage of µR constructs after treating with different ionic solutions (LiCl, NaCl, KCl, $CaCl_2$ and $MgCl_2$) and DI water (non-ionic solution).

**File Name:** Supp Video S7
**Description:** Shrinkage of µR constructs after treating with sucrose solutions, different concentration of NaCl solution and DMSO.

**File Name:** Supp Video S8
**Description:** Shrinkage of GelMA microgel constructs in 1X PBS.

**File Name:** Supp Video S9
**Description:** Shrinkage of µR constructs after treating with 1X PBS at pH 1 and 14, and culture medium at 4 degrees Celsius.

**File Name:** Supp Video S10

**Description:** Shrinkage of μR constructs in 1X and 10X PBS.

**File Name:** Supp Video S11
**Description:** Shrinkage of μR constructs with channel in 10X PBS.
